# Commonly used software tools produce conflicting and overly-optimistic AUPRC values

**DOI:** 10.1101/2024.02.02.578654

**Authors:** Wenyu Chen, Chen Miao, Zhenghao Zhang, Cathy Sin-Hang Fung, Ran Wang, Yizhen Chen, Yan Qian, Lixin Cheng, Kevin Y. Yip, Stephen Kwok-Wing Tsui, Qin Cao

## Abstract

The precision-recall curve (PRC) and the area under it (AUPRC) are useful for quantifying classification performance. They are commonly used in situations with imbalanced classes, such as cancer diagnosis and cell type annotation. We evaluated 10 popular tools for plotting PRC and computing AUPRC, which were collectively used in *>*3,000 published studies. We found the AUPRC values computed by the tools rank classifiers differently and some tools produce overly-optimistic results.

## Introduction

Many problems in computational biology can be formulated as binary classification, in which the goal is to infer whether an entity (e.g., a cell) belongs to a target class (e.g., a cell type). Accuracy, precision, sensitivity (i.e., recall), specificity, and F1 score (Supplementary Figure 1) are some of the measures commonly used to quantify classification performance, but they all require a threshold of the classification score to assign every entity to either the target class or not. The receiver operating characteristic (ROC) and precision-recall curve (PRC) avoid this problem by considering multiple thresholds [1], which allows detailed examination of the trade-off between identifying entities of the target class and wrongly including entities not of this class. It is common to summarize these curves by the area under them (AUROC and AUPRC, respectively), which is a value between 0 and 1, with a larger value corresponding to better classification performance.

When the different classes have imbalanced sizes (e.g., the target cell type has few cells), AUPRC is a more sensitive measure than AUROC [1–4], especially when there are errors among the top predictions (Supplementary Figure 2). As a result, AUPRC has been used in a variety of applications, such as reconstructing biological networks [5], identifying cancer genes [6] and essential genes [7], determining protein binding sites [8], imputing sparse experimental data [9], and predicting patient treatment response [10]. AUPRC has also been extensively used as a performance measure in benchmarking studies, such as the ones for comparing methods for analyzing differential gene expression [11], identifying gene regulatory interactions [12], and inferring cell-cell communications [13] from single-cell RNA sequencing data.

Given the importance of PRC and AUPRC, we analyzed commonly used software tools and found that they produce contrasting results, some of which are overly-optimistic.

## Results

### Basics

For each entity, a classifier outputs a score to indicate how likely it belongs to the target (i.e., “positive”) class. Depending on the classifier, the score can be discrete (e.g., random forest) or continuous (e.g., artificial neural network). Using a threshold *t*, the classification scores can be turned into binary predictions by considering all entities with a score ≥ *t* as belonging to the positive class and all other entities as not. When these predictions are compared to the actual classes of the entities, precision is defined as the proportion of entities predicted to be positive that are actually positive, while recall is defined as the proportion of actually positive entities that are predicted to be positive (Supplementary Figure 1).

The PRC is a curve that shows how precision changes with recall. In the most common way to produce the PRC, each unique classification score observed is used as a threshold to compute a pair of precision and recall values, which forms an anchor point on the PRC. Adjacent anchor points are then connected to produce the PRC.

When no two entities have the same score (Figure 1a), it is common to connect adjacent anchor points directly by a straight line [14–19] (Figure 1b). Another method uses an expectation formula, which we will explain below, to connect discrete points by piece-wise linear lines [20] (Figure 1c). The third method is to use the same expectation formula to produce a continuous curve between adjacent anchor points [17, 21] (Figure 1d). A fourth method that has gained popularity, known as Average Precision (AP), connects adjacent anchor points by step curves [15, 19, 22, 23] (Figure 1e). In all four cases, PRC estimates a function of precision in terms of recall based on the observed classification scores of the entities, and AUPRC estimates the integral of this function using trapezoids (in the direct straight line case), interpolation lines/curves (in the expectation cases), or rectangles (in the AP case).

**Figure 1.**
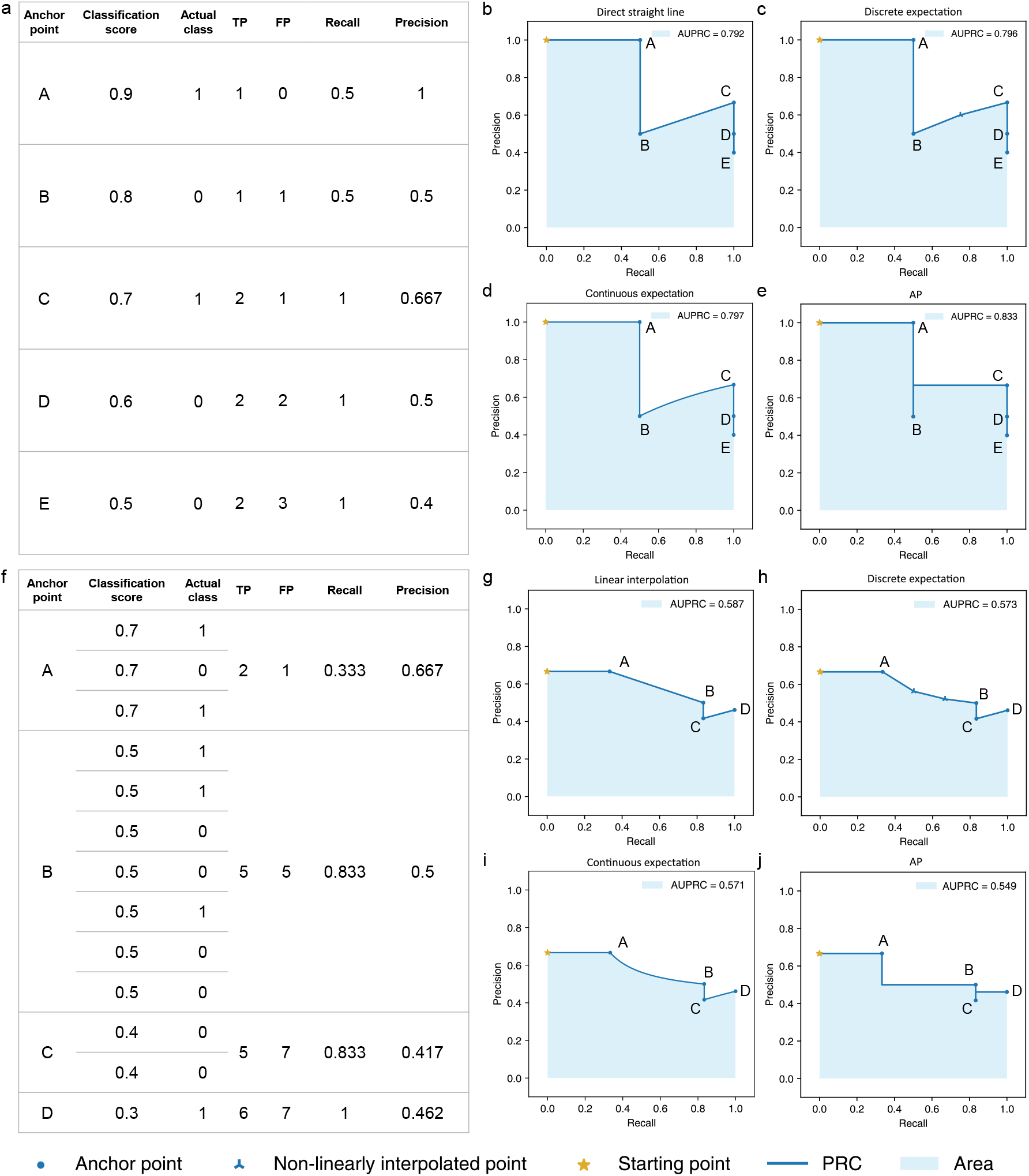
Different methods for connecting adjacent anchor points on the PRC. **a** An illustrative data set with no two entities receiving the same classification score. **b-e** Different methods for connecting adjacent anchor points when there are no ties in classification scores, namely **b** direct straight line, **c** discrete expectation, **d** continuous expectation, and **e** AP. **f** An illustrative data set with different entities receiving the same classification score. Each group of entities with the same classification score defines a single anchor point (A, B, C, and D, from 3, 7, 2, and 1 entities, respectively). **g-j** Different methods for connecting anchor point B to its previous anchor point, A, namely **g** linear interpolation, **h** discrete expectation, **i** continuous expectation, and **j** AP. In **c** and **h**, *tp* is set to 0.5 and 1 in Formula 1, respectively (Supplementary text).

When there are ties with multiple entities having the same score, which happens more easily with classifiers that produce discrete scores, these entities together define only one anchor point (Figure 1f). There are again four common methods for connecting such an anchor point to the previous anchor point, which correspond to the four methods for connecting anchor points when there are no ties (details in Supplementary text). The first method is to connect the two anchor points by a straight line [15, 18, 19] (Figure 1g). This method is known to easily produce overly-optimistic AUPRC values [2, 24], which we will explain below. The second method is to interpolate additional points between the two anchor points using a non-linear function and then connect the points by straight lines [14, 17, 20] (Figure 1h). The interpolated points appear at their expected coordinates under the assumption that all possible orders of the entities with the same score have equal probability. The third method uses the same interpolation formula as the second method but instead of creating a finite number of interpolated points, it connects the two anchor points by a continuous curve [17, 21] (Figure 1i). Finally, the fourth method comes naturally from the AP approach, which uses step curves to connect the anchor points [15, 19, 22, 23] (Figure 1j).

Using the four methods to connect anchor points when there are no ties and the four methods when there are ties can lead to very different AUPRC values (Figure 1, Supplementary Figure 3, and Supplementary text).

### Conceptual and implementation issues of some popular software tools

We analyzed 10 tools commonly used to produce PRC and AUPRC (Supplementary Table 1). Based on citations and keywords, we estimated that these tools have been used in *>*3,000 published studies in total (Methods).

The 10 tools use different methods to connect anchor points on the PRC and therefore they can produce different AUPRC values (Table 1, Supplementary Figures 4-7, and Supplementary text). As a comparison, all 10 tools can also compute AUROC, and we found most of them to produce identical values (Supplementary text).

**Table 1:**
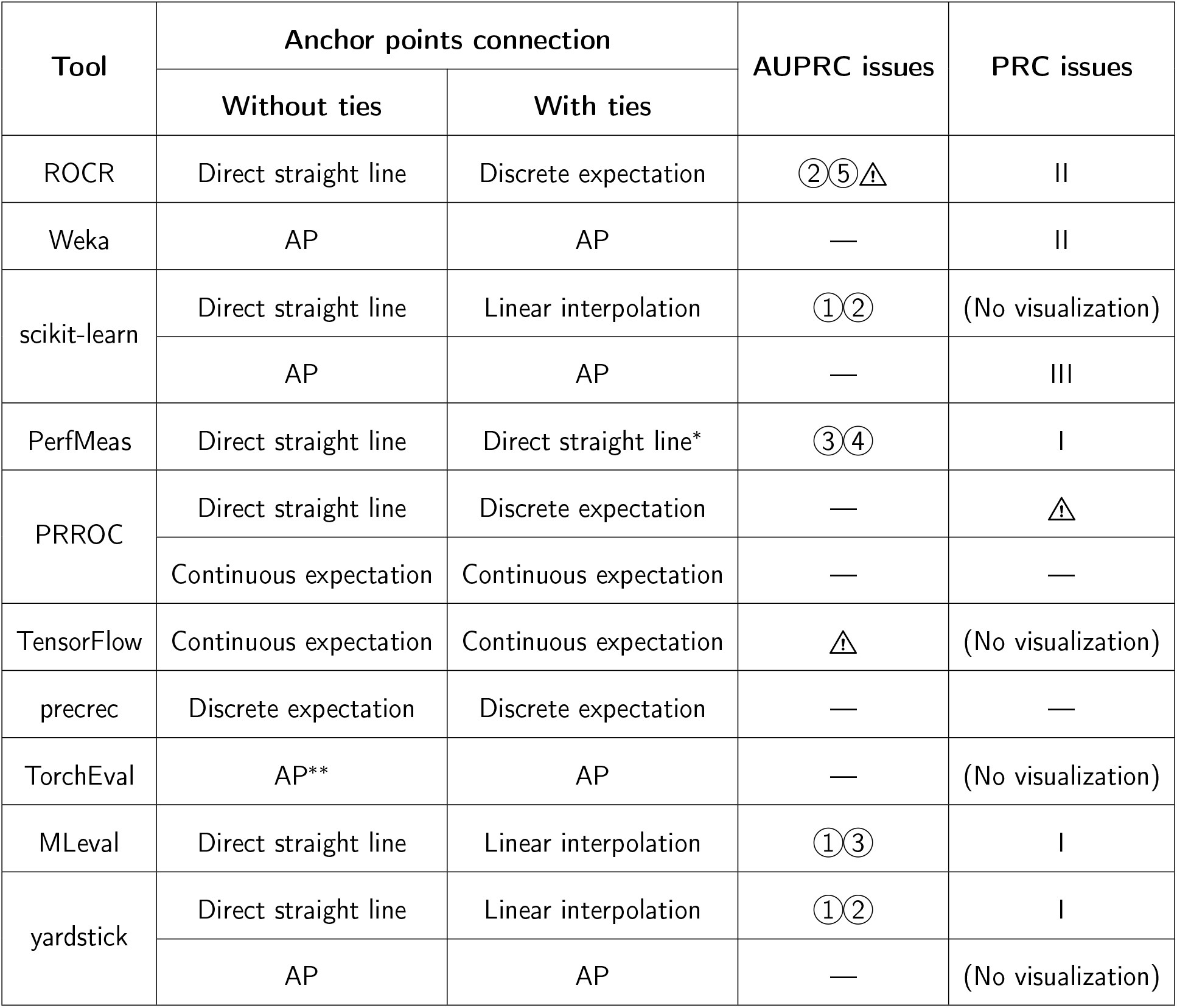
Methods used by the different software tools to connect anchor points and issues found in their calculation of AUPRC and construction of the PRC. For tools that can connect anchor points in multiple ways, we show each of them in a separate row. The AUPRC and PRC issues are defined in the text and detailed in Supplementary text. “—” means no issues found. ^*^PerfMeas orders entities with the same classification score by their order in the input and then defines anchor points as if there are no ties. ^**^The source code of TorchEval states that it uses Riemann integral to compute AUPRC, which is equivalent to AP.

| Tool | Anchor points connection |  | AUPRC issues | PRC issues |
| --- | --- | --- | --- | --- |
|  | Without ties | With ties |  |  |
| ROCR | Direct straight line | Discrete expectation | ②⑤△ | II |
| Weka | AP | AP | — | II |
| scikit-learn | Direct straight line | Linear interpolation | ①② | (No visualization) |
|  | AP | AP | — | III |
| PerfMeas | Direct straight line | Direct straight line* | ③④ | I |
| PRROC | Direct straight line | Discrete expectation | — | △ |
|  | Continuous expectation | Continuous expectation | — | — |
| TensorFlow | Continuous expectation | Continuous expectation | △ | (No visualization) |
| precrec | Discrete expectation | Discrete expectation | — | — |
| TorchEval | AP** | AP | — | (No visualization) |
| MLeval | Direct straight line | Linear interpolation | ①③ | I |
| yardstick | Direct straight line | Linear interpolation | ①② | I |
|  | AP | AP | — | (No visualization) |

We found five conceptual issues with some of these tools when computing AUPRC values (Table 1):

➀ Using the linear interpolation method to handle ties, which can produce overly-optimistic AUPRC values [2, 24]. When interpolating between two anchor points, linear interpolation produces higher AUPRC than the other three methods under conditions that can easily happen in real situations (Supplementary text)
➁ Always using (0, 1) as the starting point of the PRC (procedurally produced or conceptually derived, same for➂and ➄ below), which is inconsistent with the concepts behind the AP and non-linear expectation methods when the first anchor point with a non-zero recall does not have a precision of one (Supplementary text)
➂ Not producing a complete PRC that covers the full range of recall values from zero to one
➃ Ordering entities with the same classification score by their order in the input and then handling them as if they have distinct classification scores
➄ Not putting all anchor points on the PRC

These issues can lead to overly-optimistic AUPRC values or change the order of two AUPRC values (Supplementary text, Supplementary Figures 8-13).

Some of these tools also produce a visualization of the PRC. We found three types of issues with these visualizations (Table 1):

I. Producing a visualization of PRC that has the same issue(s) as in the calculation of AUPRC
II. Producing a PRC visualization that does not always start the curve at a point with zero recall
III. Producing a PRC visualization that always starts at (0, 1)

Finally, we also found some programming bugs and noticed that some tools require special attention for correct usage (both marked by 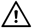 in Table 1).

### Inconsistent AUPRC values and contrasting classifier ranks produced by the popular tools

To see how the use of different methods by the 10 tools and their other issues affect PRC analysis in practice, we applied them to evaluate classifiers in four realistic scenarios.

In the first scenario, we analyzed data from a COVID-19 study [25] in which patient blood samples were subjected to Cellular Indexing of Transcriptomes and Epitopes by Sequencing (CITE-seq) assays [26]. We constructed a classifier for predicting CD4^+^ T cells, which groups the cells based on their transcriptome data alone and assigns a single cell type label to each group. Using cell type labels defined by the original authors as reference, which were obtained using both antibody-derived tags (ADTs) and transcriptome data, we computed the AUPRC of the classifier. Figure 2a shows that the 10 tools produced 6 different AUPRC values, ranging from 0.416 to 0.684. In line with the conceptual discussions above, the AP method generally produced the smallest AUPRC values while the linear interpolation method generally produced the largest, although individual issues of the tools created additional variations of the AUPRC values computed.

**Figure 2.**
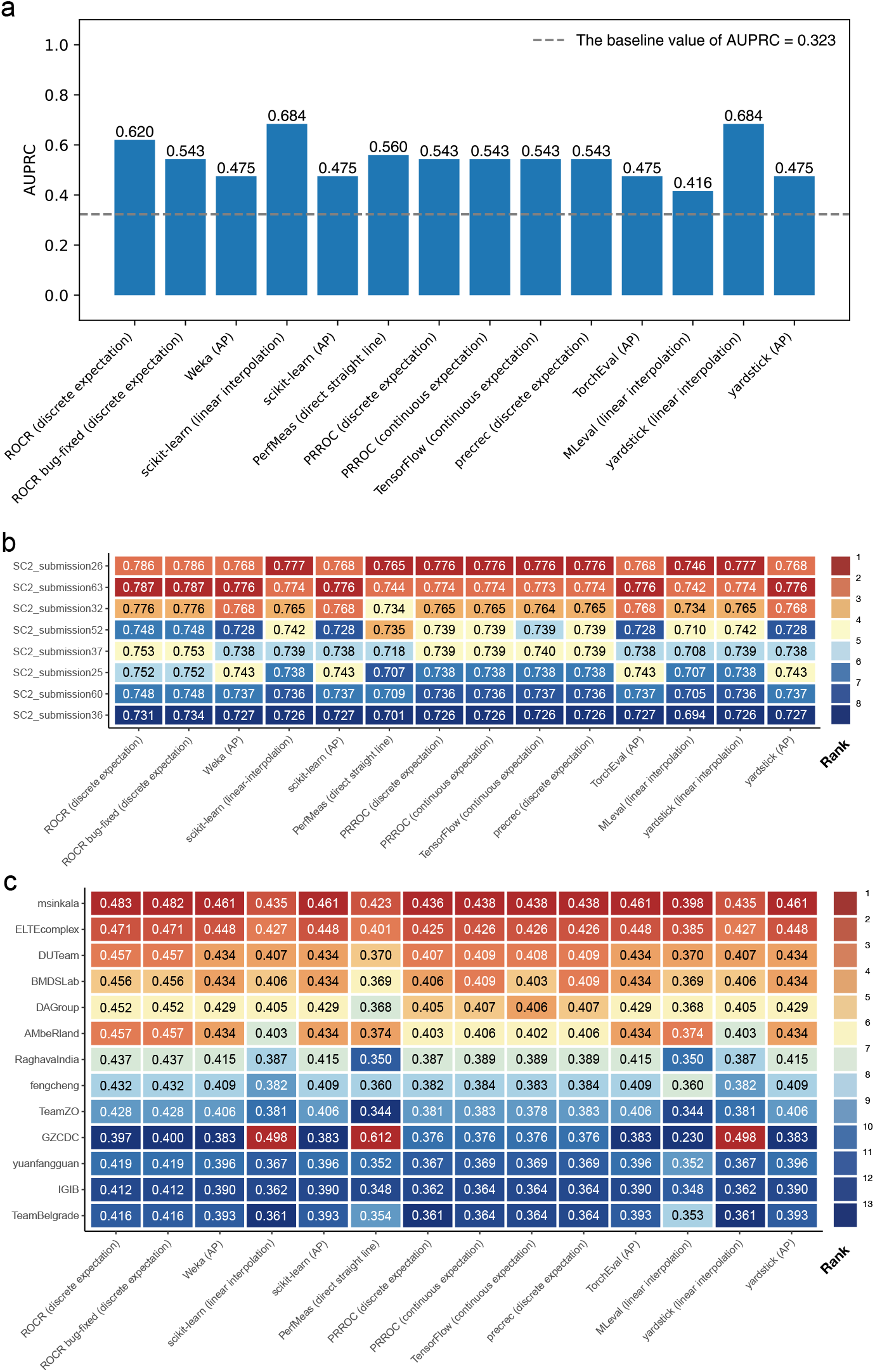
The AUPRC values computed by the 10 tools in several realistic scenarios. **a** Predicting CD4^+^ T cells from single-cell transcriptomic data. **b** Predicting inflammatory bowel disease cases that belong to the Ulcerative Colitis subtype in the sbv IMPROVER Metagenomics Diagnosis for Inflammatory Bowel Disease Challenge. Only the top 8 submissions according to PRROC (discrete expectation) AUPRC values are included. **c** Predicting cases with preterm prelabor rupture of membranes in the DREAM Preterm Birth Prediction Challenge. In **b** and **c**, each entry shows the AUPRC value and the background color indicates its rank among the competitors.

In the second scenario, we compared the performance of different classifiers that predict whether a patient has the Ulcerative Colitis (UC) subtype of inflammatory bowel disease (IBD) or does not have IBD, based on metagenomic data (processed taxonomy-based profile) [27]. The predictions made by these classifiers were submitted to the sbv IMPROVER Metagenomics Diagnosis for IBD Challenge. Their performance was determined by comparing against diagnosis of these patients based on clinical, endoscopic, and histological criteria. Figure 2b shows that based on the AUPRC values computed, the 10 tools ranked the classifiers differently. For example, among the top 8 submissions with the highest performance, the classifier in submission 26 was ranked first in 8 cases, sole second place in 2 cases, and tied second place with another classifier in 4 cases (Figure 2b and Supplementary Figure 14). We observed similar rank flips when considering the top 30 submissions (Supplementary Figures 15 and 16).

In the third scenario, we compared the performance of different classifiers in identifying preterm prelabor rupture of the membranes (PPROM) cases from normal pregnancy in the DREAM Preterm Birth Prediction Challenge [28]. Based on the AUPRC values produced by the 10 tools, the 13 participating teams were ranked very differently (Figure 2c and Supplementary Figure 17). For example, Team “GZCDC” was ranked first (i.e, highest) in 3 cases, tenth in 4 cases, and thirteenth (i.e., lowest) in 7 cases. In addition to differences in the ranks, some of the AUPRC values themselves are also very different. For example, the AUPRC values computed by PerfMeas and MLeval have a Pearson correlation of -0.759, which shows that their evaluations of the 13 teams were almost completely opposite.

In the fourth scenario, we compared 29 classifiers that predicted target genes of transcription factors in the DREAM5 challenge [29]. Again, some classifiers received very different ranking based on the AUPRC values computed by the different tools (Supplementary Figures 18 and 19). For example, the classifier named “Other4” was ranked second based on the AUPRC values computed by PerfMeas but it was ranked twenty-fifth based on the AUPRC values computed by MLeval. In general, tools that use the discrete expectation, continuous expectation, and AP methods are in good agreements in this scenario, but they differ substantially from tools that use the linear interpolation method.

## Conclusions

Due to their highly technical nature, it is easy to overlook the inconsistencies and issues of the software tools used for producing PRC and AUPRC. Some possible consequences include reporting overly-optimistic AUPRC, ranking classifiers differently by different tools, and introducing biases to the evaluation process, such as inflating the AUPRC of classifiers that produce discrete scores.

## Methods

### Information about the tools

In this study, we included 12 tools commonly used for PRC and ROC analyses (Supplementary Table 1). For each tool, we analyzed the latest stable version of it as of August 15, 2023. Because TorchEval had not released a stable version, we analyzed the latest version of it, version 0.0.6. Among the 12 tools, ten can compute both AUROC and AUPRC, while the remaining two can only compute AUROC. We focused on these 10 tools in the study of PRC and AUPRC. Some tools provide multiple methods for computing AUROC/AUPRC.

For tools with an associated publication, we obtained its citation count from Google Scholar. If a tool has multiple associated publications, we selected the one with the largest number of citations. As a result, the citation counts we report in Supplementary Table 1 are underestimates if different publications associated with the same tool are not always cited together.

The Comprehensive R Archive Network (CRAN) packages PerfMeas and MLeval did not have an associated formal publication but only release notes. In each of these cases, we used the package name as keyword to search on Google Scholar and then manually checked the publications returned to determine the number of publications that cited these packages.

The CRAN package yardstick also did not have an associated formal publication. However, we were not able to use the same strategy as PerMeas and MLeval to determine the number of publications that cited the yardstick package since “yardstick” is an English word and the search returned too many publications to be verified manually. Therefore, we only counted the number of publications that cited yardstick’s release note, which is likely an underestimate of the number of publications that cited yardstick.

All citation counts were collected on October 9, 2023.

For tools with an associated formal publication, based on our collected lists of publications citing the tools, we further estimated the number of times the tools were actually used in the studies by performing keyword-based filtering. Specifically, if the main text or figure captions of a publication contains either one of the keywords “AUC” and “AUROC”, we assumed that the tool was used in that published study to perform ROC analysis. In the case of PRC, we performed filtering in two different ways and reported both sets of results in Supplementary Table 1. In the first way, we assumed a tool was used in a published study if the main text or figure captions of the publication contains any one of the following keywords: “AUPR”, “AU-PR”, “AUPRC”, “AU-PRC”, “AUCPR”, “AUC-PR”, “PRAUC”, “PR-AUC”, “area under the precision recall”, and “area under precision recall”. In the second way, we assumed a tool was used in a published study if the main text or figure captions of the publication contains both “area under” and “precision recall”.

For the CRAN packages PerfMeas and MLeval, we estimated the number of published studies that actually used them by searching Google Scholar using the above three keyword sets each with the package name appended. We found that for all the publications we considered as using the packages in this way, they were also on our lists of publications that cite these packages. We used the same strategy to identify published studies that used the CRAN package yardstick. We found that some of these publications were not on our original list of publications that cite yardstick, and therefore we added them to the list and updated the citation count accordingly.

TorchEval was officially embedded into PyTorch in 2022. Due to its short history, among the publications that cite the PyTorch publication, we could not find any of them that used the TorchEval library.

## Data collection and processing

We used four realistic scenarios to illustrate the issues of the AUPRC calculations.

In the first scenario, we downloaded CITE-seq data produced from COVID-19 patient blood samples by the COVID-19 Multi-Omic Blood ATlat (COMBAT) consortium [25]. We downloaded the data from https://doi.org/10.5281/zenodo.6120249 and used the data in the “COMBAT-CITESeq-DATA” archive in this study. We then used a standard procedure to cluster the cells based on the transcriptome data and identified CD4^+^ T cells. Specifically, we extracted the raw count matrix of the transcriptome data and ADT features (“X” object) and the annotation data frame (“obs” object) from the H5AD file. We dropped all ADT features (features with names starting with “AB-”) and put the transcriptome data along with the annotation data frame into Seurat (version 4.1.1). We then log-normalized the transcriptome data (method “NormalizeData()”, default parameters), identified highly-variable genes (method “FindVariableFeatures()”, number of variable genes set to 10,000), scaled the data (method “ScaleData()”, default parameters), performed principal component analysis (method “RunPCA()”, number of principal components set to 50), constructed the shared/k-nearest neighbor (SNN/kNN) graph (method “FindNeighbours()”, default parameters), and performed Louvain clustering of the cells (method “FindClusters()”, default parameters). We then extracted the clustering labels generated and concatenated them with cell type, major subtype, and minor subtype annotations provided by the original authors, which were manually curated using both ADT and transcriptome information.

Our procedure produced 29 clusters, which contained 836,148 cells in total. To mimic a classifier that predicts CD4^+^ T cells using the transcriptome data alone, we selected one cluster and “predicted” all cells in it as CD4^+^ T cells and all cells in the other 28 clusters as not, based on which we computed an AUPRC value by comparing these “predictions” with the original authors’ annotations. We repeated this process for each of the 29 clusters in turn, and chose the one that gave the highest AUPRC as the final cluster of predicted CD4^+^ T cells.

For the second scenario, we obtained the data set used in the sbv IMPROVER (Systems Biology Verification combined with Industrial Methodology for PROcess VErification in Research) challenge on inflammatory bowel disease diagnosis based on metagenomics data [27]. The challenge involved 12 different tasks, and we focused on the task of identifying UC samples from non-IBD samples using the processed taxonomy-based profile as features. The data set contained 32 UC samples and 42 non-IBD samples, and therefore the baseline AUPRC was 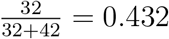. There were 60 submissions in total, which used a variety of classifiers. We obtained the classification scores in the submissions from Supplementary Information 4 of the original publication [27]. When we extracted the classification scores of each submission, we put the actual positive entities before the actual negative entities. This ordering did not affect the AUPRC calculations of most tools except those of PerfMeas, which depend on the input order of the entities with the same classification score.

To see how the different tools rank the top submissions, we first computed the AUPRC of each submission using PRROC (option that uses the discrete expectation method to handle ties) since we did not find any issues with its AUPRC calculations (Table 1). We then analyzed the AUPRC values produced by the 10 tools based on either the top 8 (Figure 2b and Supplementary Figure 14) or top 30 (Supplementary Figures 15 and 16) submissions.

For the third scenario, we downloaded the data set used in the Dialogue on Reverse Engineering Assessment and Methods (DREAM) Preterm Birth Prediction Challenge [28] from https://www.synapse.org/#!Synapse:syn22127152. We collected the classification scores, from the object “prpile” in each team’s RData file, and the actual classes produced based on clinical evidence, from “anoSC2_v21_withkey.RData” (https://www.synapse.org/#!Synapse:syn22127343). The challenge contained 7 scenarios, each of which had 2 binary classification tasks. For each scenario, 10 different partitioning of the data into training and testing sets were provided. We focused on the task of identifying PPROM cases from the controls under the D2 scenario defined by the challenge. For this task, the baseline AUPRC value averaged across the 10 testing sets was 0.386. There were 13 participating teams in total. For each team, we extracted its classification scores and placed the actual positive entities before the actual negative entities. For submissions that contained negative classification scores, we re-scaled all the scores to the range between 0 and 1 without changing their order since TensorFlow expects all classification scores to be between zero and one (Supplementary text). Finally, for each team, we computed its AUPRC using each of the 10 testing sets and reported their average. We note that the results we obtained by using PRROC (option that uses the continuous expectation method to handle ties) were identical to those reported by the challenge organizer.

For the fourth scenario, we obtained the data set used in the DREAM5 challenge on reconstructing transcription factor-target networks based on gene expression data [29]. The challenge included multiple networks and we focused on the *E. coli in silico* Network 1, which has a structure that corresponds to the real E. coli transcriptional regulatory network [29]. We obtained the data from Supplementary Data of the original publication [29]. There were 29 submissions in total. For each submission, we extracted the classification scores of the predicted node pairs (each pair involves one potential transcription factor and one gene it potentially regulates) from Supplementary Data 4 and compared them with the actual classes (positive if the transcription factor actually regulates the gene; negative if not) in the gold-standard network from Supplementary Data 3. Both the submissions and the gold-standard were not required to include all node pairs. To handle this, we excluded all node pairs in a submission that were not included in the gold-standard (because we could not judge whether they are actual positives or actual negatives), and assigned a classification score of 0 to all node pairs in the gold-standard that were not included in a submission (because the submission did not give a classification score to them). The gold-standard contained 4,012 interacting node pairs and 274,380 non-interacting node pairs, and therefore the baseline AUPRC value was 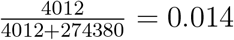.

## Data availability

All data used in this study can be accessed following the procedure described in the “Data collection and processing” part of the Methods section.

## Code availability

We have submitted all the code used in our study for the review process. We will create a GitHub repository to allow public free access to our code.

Acknowledgments

QC is supported by National Natural Science Foundation of China under Award Number 32100515. KYY is supported by National Cancer Institute of the National Institutes of Health under Award Number P30CA030199, National Institute on Aging of the National Institutes of Health under Award Numbers U54AG079758 and R01AG085498, and internal grants of Sanford Burnham Prebys Medical Discovery Institute. The content is solely the responsibility of the authors and does not necessarily represent the official views of the National Institutes of Health.

## Author contributions

KYY, SKWT and QC conceived and supervised the project. WC, CM, KYY and QC designed the computational experiments and data analyses. WC, CM and YC surveyed and collected the tools. WC, CM, ZZ, CSHF and RW prepared the data. WC and CM conducted the computational experiments. WC, CM, ZZ and RW performed the data analyses. All the authors interpreted the results. WC, CM, KYY and QC wrote the manuscript.

## Supplementary text

### Mathematical details of PRC and AUPRC

#### The non-linear interpolation formula

Suppose *A* is an anchor point and *B* is its next anchor point, defined by score thresholds *t*_*A*_ and *t*_*B*_, respectively. In the most standard way to define anchor points, both *t*_*A*_ and *t*_*B*_ are classification scores of some entities, and *t*_*B*_ is the largest classification score of the entities that is smaller than *t*_*A*_. If we predict all entities with a classification score ≥*t*_*A*_ to belong to the positive class, there are *TP*_*A*_ true positives and *FP*_*A*_ false positives. Similarly, if we predict all entities with a classification score *≥ t*_*B*_ to belong to the positive class, there are *TP*_*B*_ true positives and *FP*_*B*_ false positives, where *TP*_*B*_ *≥ TP*_*A*_ and *FP*_*B*_ *≥ FP*_*A*_. The value *m* = (*TP*_*B*_ − *TP*_*A*_) + (*FP*_*B*_− *FP*_*A*_) is the number of entities with a classification score of *t*_*B*_. If *m >* 1, the interpolation methods interpolate additional point(s) between *A* and *B*.

One way to interpolate these additional points is to place them at their expected locations, assuming that all possible orders of the *m* entities have the same probability. Specifically, suppose at an additional point, there are *TP*_*A*_ + *tp* true positives, where *tp* is an integer between 1 and *TP*_*B*_ − *TP*_*A*_ − 1. The expected number of false positives at this point is 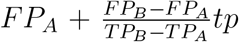. As a result, the x and y coordinates of this additional point, which respectively correspond to its recall and precision, are:

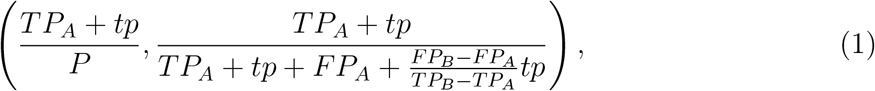

where *P* is the total number of actual positives.

The above method, proposed by Davis and Goadrich [2], interpolates *TP*_*B*_ − *TP*_*A*_− 1 points with equal spacing along the x-axis between *A* and *B*. We refer to it as the “discrete expectation” method (Figure 1h and Supplementary Figure 3h). It should be noted that if we set *tp* to *TP*_*B*_ − *TP*_*A*_, the formula produces the coordinates of anchor point *B* exactly. This is important for the continuous expectation method to be discussed below.

From Formula 1, it can be seen that precision does not necessarily change linearly with recall [2].

In the literature, some small variations of this method have been proposed. They also use Formula 1 to determine coordinates of the interpolated points but instead of placing *TP*_*B*_ − *TP*_*A*_ − 1 points with equal spacing along the x-axis between *A* and *B*, they place the points at fixed intervals of false positive count [30] or recall [20] by setting *tp* to corresponding (possibly non-integer) values accordingly. We call all these variations “discrete expectation” for simplicity. Also, some tools use the discrete expectation method to produce additional points even when there are no ties in classification scores (Figure 1c and Supplementary Figure 3c).

In Formula 1, if instead of considering just a finite number of discrete values of *tp*, it is allowed to take any real value between 0 and *TP*_*B*_ − *TP*_*A*_, anchor points *A* and *B* will be connected by a continuous curve [30, 31]. We refer to this method as the “continuous expectation” method (Figure 1i and Supplementary Figure 3i). The area under the PRC between *A* and *B* can be computed by integrating the function defined by Formula 1:

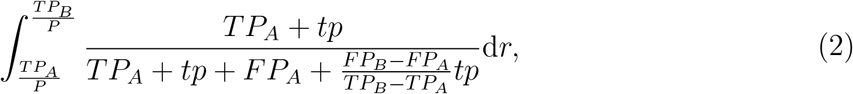

where 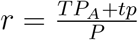.

Let 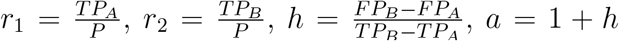, and 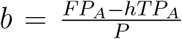, the integral in Formula 2 becomes [30]:

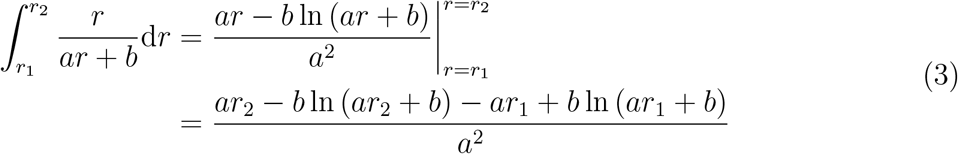

A previous study showed that practically, the discrete expectation and continuous expectation methods produce comparable results across various binary classification scenarios [30].

### The Average Precision method

The Average Precision (AP) method [32] connects anchor points by step curves (Figure 1e,j and Supplementary Figure 3e,j). The AUPRC computed by the AP method represents the weighted mean of precision at different anchor points, where the weight is the increase in recall from the previous anchor point [15]:

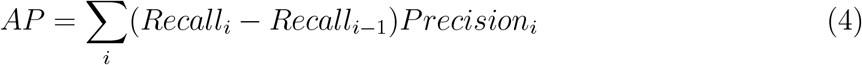

In the formula, the summation is taken over all anchor points *i*.

In information retrieval, where AP is widely applied, there is another well-accepted definition of AP. In this definition, instead of computing average precision at the anchor points defined by the unique classification scores of the entities, it computes average precision at a fixed number of points evenly spaced along the x-axis [33–35].

In computer vision, AP has become one of the most common assessment metrics in image segmentation. A recent study reported that five different definitions of AP coexist in bioimage segmentation analysis [36].

The tools we surveyed in the current study only implemented AP as defined by Formula 4 but not any of these other variations.

### The starting point of PRC

In the main text, we defined the anchor points by the unique classification scores that the entities receive from the classifier. Specifically, for each unique score *t*, an anchor point is defined, which predicts all entities with a score *≥ t* as positive and all other entities as not. If among the entities receiving the highest score at least one of them actually belongs to the positive class, the first anchor point will have a non-zero recall. In order for the PRC to cover the whole range of recall values from zero to one, we need a way to define a starting point that always has zero recall.

If the AP method is used to connect the starting point and the first anchor point, we can simply set the starting point to (0, *Precision*_1_), where *Precision*_1_ is the precision of the first anchor point. This is obvious if the first anchor point does not involve any entities actually coming from the positive class, in which case the recall of it would be zero and thus the starting point is identical to the first anchor point. On the other hand, if the first anchor point involves at least one entity actually coming from the positive class, by setting the precision of the starting point to be the same as the first anchor point, in the PRC these two points will be connected by a horizontal line, which matches the definition of AP in Formula 4.

If the continuous expectation method is used to connect the starting point and the first anchor point, we can use Formula 1 to determine the coordinates of the points on this connection curve. Substituting both *TP*_*A*_ and *FP*_*A*_ by 0, *TP*_*B*_ by *TP*_1_ (number of true positives at the first anchor point), and *FP*_*B*_ by *FP*_1_ (number of false positives at the first anchor point), the coordinates of the connection points become:

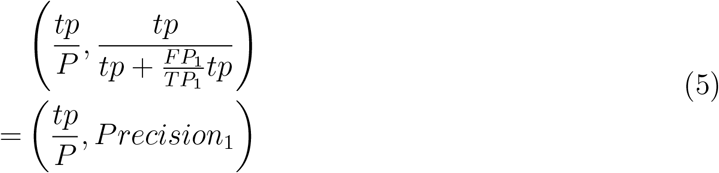

As *tp* approaches 0, the coordinates approach (0, *Precision*_1_).

Therefore, for both the AP method and the continuous expectation method, the starting point of the PRC is (0, *Precision*_1_).

Since the discrete expectation and continuous expectation methods both use the same way to determine locations of the interpolated points, we argue that when the discrete expectation method is used, it is reasonable to also set the starting of the PRC to (0, *Precision*_1_) following the continuous expectation method. In contrast, for the linear interpolation method, we were not able to find in the literature any proposed way to determine its starting point with a sound justification.

### The baseline value of AUPRC

It is useful to benchmark the AUPRC value of a classifier against that of a baseline classifier. A commonly used baseline classifier is one that gives the same score to every entity. The corresponding PRC contains only one anchor point at 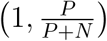, where *P* and *N* are the actual numbers of positives and negatives, respectively.

If the AP method or the continuous expectation method is used to compute AUPRC, as mentioned above, the PRC of the baseline classifier will have a starting point of 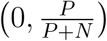, and thus the PRC is simply a horizontal line with an AUPRC of 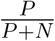. This quantity is called the baseline AUPRC value, which is simply the fraction of actual positives among all the entities [17, 37].

In the case of continuous expectation, we can also directly compute the AUPRC using Formula 3 by substituting *TP*_*A*_ and *FP*_*A*_ by 0, *TP*_*B*_ by *P*, and *FP*_*B*_ by *N*. Accordingly, the variables used in the formula will take the following values:

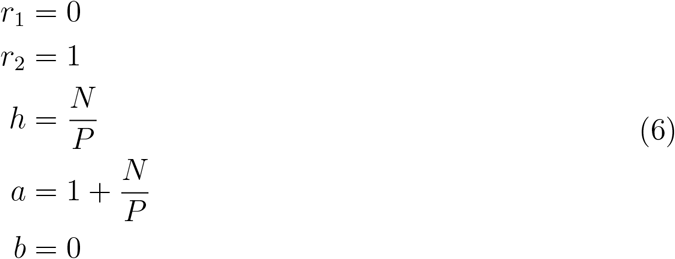

Formula 3 then becomes:

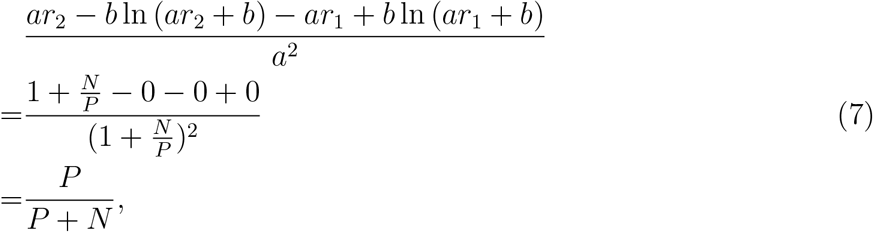

which is the same as the value we obtained by considering the starting point of the PRC.

### The five issues of deriving PRC and AUPRC

#### Description of the five issues

We found five issues when using the tools we analyzed to compute AUPRC values. These issues can be most easily explained by considering the corresponding PRC (Supplementary Figures 4-7), no matter the tools indeed first determine the points on the PRC and then compute the area under it accordingly, or rather compute AUPRC directly without first determining the points.

➀Linear interpolation: When the linear interpolation method is used to handle ties in classification scores, theoretically the resulting AUPRC can be either larger (Figure 1g-j) or smaller (Supplementary Figure 3g-j) than the AUPRC computed by the other three methods. Specifically, the linear interpolation method produces a larger AUPRC between the two anchor points than the AP method if the tie-associated anchor point (point *B* in Figure 1g,j) has a smaller precision than the previous anchor point (point *A* in the figures). Conversely, the linear interpolation method produces a smaller AUPRC between the two anchor points than the AP method if the tie-associated anchor point has a larger precision than the previous anchor point (Supplementary Figure 3g,j).

To compare the AUPRC values between the two anchor points produced by the linear interpolation and continuous expectation methods, we check whether the curve produced by the latter is convex or concave. Let 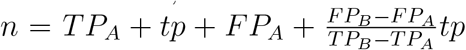, then according to Formula 1,

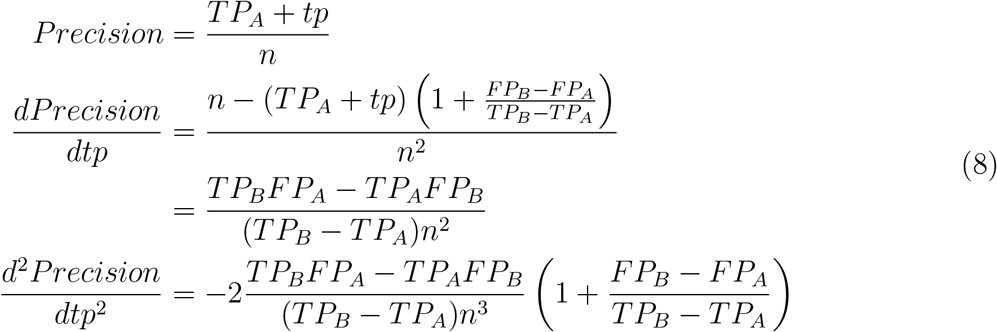

Since *TP*_*B*_ *≥ TP*_*A*_ and *FP*_*B*_ *≥ FP*_*A*_, the sign of 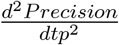 depends only on *TP*_*B*_ *FP*_*A*_ − *TP*_*A*_*FP*_*B*_. There are three possibilities:

1. If *FP*_*B*_ = *FP*_*A*_ =0 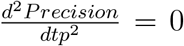. This means the continuous expectation method also connects the two anchor points by a straight line. Therefore the AUPRC between the two anchor points computed by the linear interpolation method is the same as the one computed by the continuous expectation method.
2. If *FP*_*A*_ = 0 but, 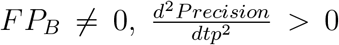. This means the continuous expectation method connects the two anchor points by a convex curve. Therefore the AURPC between the two anchor points computed by the linear interpolation method is larger than the one computed by the continuous expectation method.
3. If *FP*_*A*_ *≠* 0 and, 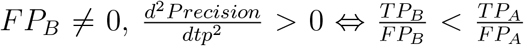. This means the continuous expectation method connects the two anchor points by i) a convex curve if the TP-to-FP ratio decreases from anchor point *A* to anchor point *B*, ii) a straight line if the TP-to-FP ratio does not change from *A* to *B*, and iii) a concave curve if the TP-to-FP ratio increases from *A* to *B*. Accordingly, the linear interpolation method produces a larger, equal, and smaller AUPRC between the two anchor points than the continuous expectation method, respectively.

This analysis also applies to the comparison between the linear interpolation method and the discrete expectation method, because the points interpolated by the discrete expectation method are all on the continuous curve interpolated by the continuous expectation method. In practice, classifiers usually have a general decreasing trend of precision (and TP-to-FP ratio) as recall increases [38] because the entities receiving the highest classification scores are usually the ones that the classifier is most confident to be positives. Therefore, the linear interpolation method tends to be overly-optimistic as compared to the other three methods for connecting anchor points when multiple entities receive the same classification score.

➁ Using (0, 1) as the starting point: As explained above, for both the continuous expectation and AP methods, the starting point of the PRC should be (0, *Precision*_1_), where *Precision*_1_ is the precision of the first anchor point. When *Precision*_1_ is not equal to one, setting the starting point to (0, 1) will inflate the resulting AUPRC.
➂ Incomplete PRC: The PRC is meant to show how the precision changes with recall over the whole range of possible values of recall (from 0 to 1). Some tools do not achieve it because they do not connect a point with zero recall to the first anchor point when the first anchor point has a non-zero recall.
➃ Arbitrary tie breaking: When there are *x* entities with the same classification score, instead of generating one anchor point, *x* anchor points are generated as if the entities all have different classification scores. The order of these entities follow their input order. As a result, running a tool with such an issue multiple times with different input orders of these entities can produce different AUPRC values. Consequently, the ranking of different classifiers can be affected by the input order of the entities in ties, instead of depending on their performance alone.
➄ Omitting anchor points: When there are multiple anchor points with the same recall, only the point with the highest precision is kept, which changes the resulting AUPRC value.

#### Consequences of the five issues

Among the five issues, four of them (➀, ➁, ➃, and ➄) can lead to overly-optimistic AUPRC values when there are ties in classification scores (Supplementary Table 2). Issue 5 (omitting anchor points) can also lead to overly-optimistic AUPRC values even when there are no ties in classification scores. Supplementary Figure 8 provides some illustrative examples.

In addition, all five issues can change the order of two AUPRC values, which means a better-performing classifier can receive a lower AUPRC than a worse-performing classifier due to the issues (Supplementary Table 2, Supplementary Figures 9-13).

### Detailed discussions of individual tools

#### ROCR

ROCR [14] is a tool in the R software environment for visualizing the performance of classifiers (Supplementary Table 1). We used the performance() function of ROCR to construct PRC and compute AUPRC. When there are no ties, ROCR connects adjacent anchor points by a straight line; When there are ties, it uses the discrete expectation method to connect adjacent anchor points to compute AUPRC (Table 1, Supplementary Figures 4a, 5a, 6a, and 7a). Specifically, ROCR uses the expectation formula (Formula 1) to interpolate additional points with a step size of *tp* = 1 when two adjacent anchor points differs in their true positive counts by more than 2.

Regarding the calculation of AUPRC, we found that ROCR always starts the PRC at (0, 1) (i.e., Issue ➁). In addition, when there are consecutive anchor points with the same recall, ROCR retains only the one with the highest precision (i.e., Issue ➄).

Regarding the PRC visualization, we found that ROCR only connects anchor points but not the starting point at zero recall (i.e., Issue II), which is inconsistent with the AUPRC value it produces.

In addition, we found a programming bug (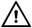) in the source code of ROCR that can lead to AUPRC values larger than one. As an example, we ran ROCR on a data set using the following code:

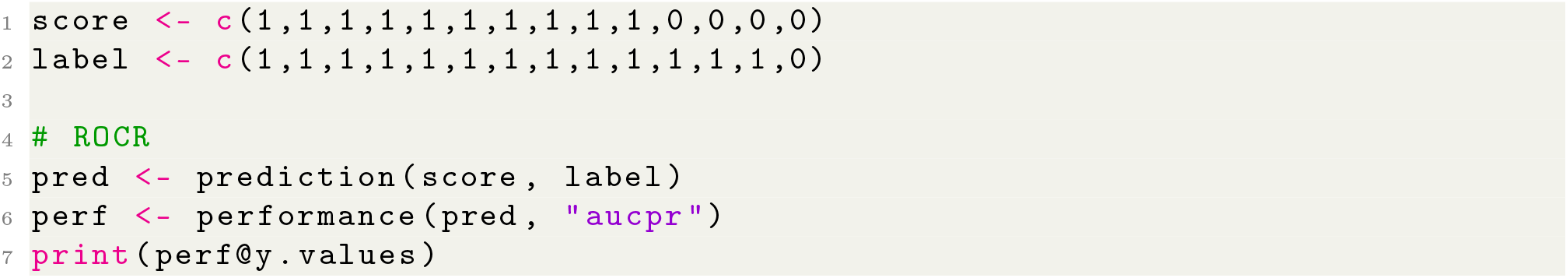

ROCR returned an AUPRC value of 1.003. We found that the problem is caused by incorrect indices of the points on the PRC. Specifically, when additional points are interpolated between two anchor points, they are added to the list of points and therefore the indices of all points after them should be shifted. However, ROCR wrongly uses their original indices. The code from ROCR’s GitHub repository (https://github.com/cran/ROCR/blob/master/R/performance_measures.R) is listed as follows:

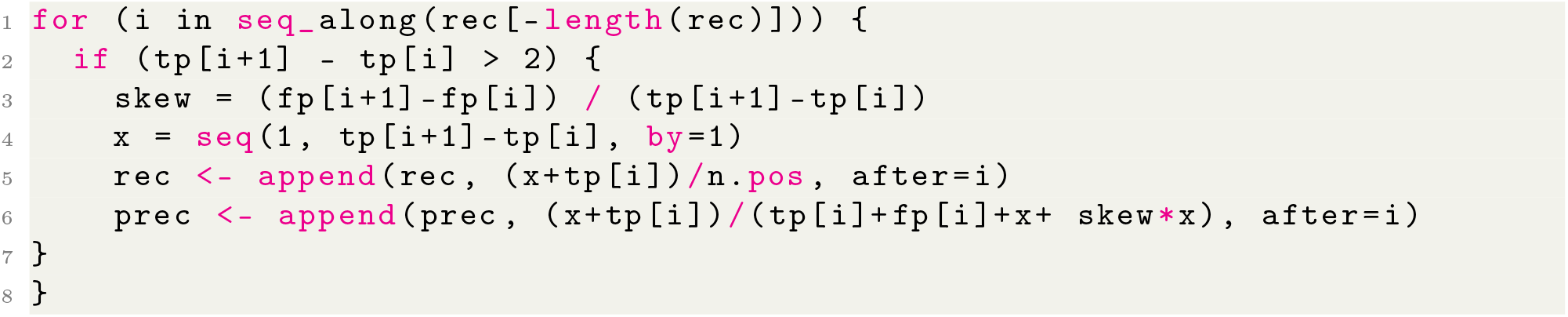

We fixed this bug by correcting the indices of the precision and recall lists after each interpolation:

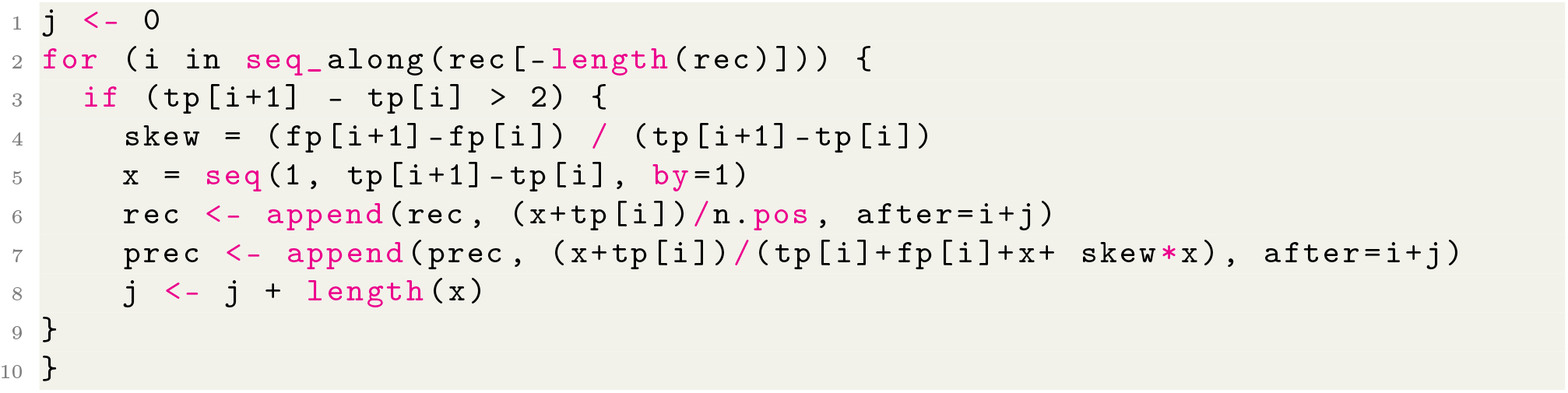

After fixing the bug, the resulting AUPRC of the example is 0.991. In the main text, all results and discussions about ROCR are about this bug-fixed version unless otherwise stated. We did not make any additional changes in response to the other issues of ROCR.

#### Weka

Weka (Waikato Environment for Knowledge Analysis) [22] is a Java-based tool for machine learning and data mining (Supplementary Table 1). It provides AUPRC calculations and PRC visualization. We used the graphical user interface of Weka to construct PRC and compute AUPRC. Underlying, the former uses the getCurve() method of the weka.classifiers.evaluation.ThresholdCurve class, and the latter uses the getPRCArea() method of the same class. Weka uses the AP method to compute AUPRC (Table 1, Supplementary Figures 4b, 5b, 6b, and 7b). In the PRC visualization, it only connects the anchor points and therefore does not start the PRC at zero recall when the first anchor point has a non-zero recall (i.e., Issue II).

#### scikit-learn

scikit-learn [15] is a Python library that provides efficient tools for machine learning and statistical modeling (Supplementary Table 1). It provides two options for computing AUPRC (Table 1), by the direct straight line/linear interpolation method (Supplementary Figures 4c, 5c, 6c, and 7c), and the AP method (Supplementary Figures 4d, 5d, 6d, and 7d), respectively. For the first option, we used the precision_recall_curve() and auc() functions of the sklearn.metrics module to compute AUPRC. For the second option, we used the Precision-RecallDisplay.from_predictions() function of the skleran.metrics module to construct PRC and the average_precision_score() function of the same module to compute AUPRC.

For the first option, if the *sklearn*.*metrics*.*precision_recall_curve* function is used to construct the PRC and then the *sklearn*.*metrics*.*auc* function is used to compute AUPRC, the linear interpolation method is used (i.e., Issue ➀) and the PRC always starts at (0, 1) (i.e.,Issue ➁).

On the manual pages of scikit-learn (based on the version in August 2023), some cautionary notes are provided regarding its AUPRC calculations:

- On the manual page about its model evaluation modules (https://scikit-learn.org/stable/modules/model_evaluation.html#precision-recall-f-measure-metrics), it is mentioned that the linear interpolation method can lead to overly-optimistic AUPRC values: “References [Davis2006] and [Flach2015] describe why a linear interpolation of points on the precision-recall curve provides an overly-optimistic measure of classifier performance. This linear interpolation is used when computing area under the curve with the trapezoidal rule in auc.” Despite this cautionary note, some recently published studies still used this method to compute AUPRC [39–44].
- On the manual page about the AP method, it is mentioned that the resulting AUPRC is different from the one computed by connecting adjacent anchor points by direct straight lines: “This implementation is not interpolated and is different from computing the area under the precision-recall curve with the trapezoidal rule, which uses linear interpolation and can be too optimistic.”

For the second option, the PRC visualization provided by scikit-learn always starts at (0, 1) (i.e., Issue III). We also noticed that in earlier versions of scikit-learn (before version 0.19), the second option of scikit-learn uses direct straight lines to connect anchor points, which is inconsistent with the AP approach meant to be taken by this option.

#### PerfMeas

PerfMeas (Performance Measures) [16] is an R library for computing performance measures of classification results (Supplementary Table 1). It provides AUPRC calculations and PRC visualizations (Table 1, Supplementary Figures 4e, 5e, 6e, and 7e). We used the precision.recall.curves.plot() function to construct PRC, and the precision.at.all.recall.levels() and trap.rule.integral() functions to compute AUPRC. PerfMeas uses direct straight lines to connect adjacent anchor points no matter whether there are ties in classification scores (i.e., Issue ➀) or not.

In the calculation of AUPRC, PerfMeas only computes the area under the points it defines (i.e., Issue ➂), including the anchor points that correspond to the unique classification scores and additional points produced by following the input order of entities with the same classification score and handling them as if there are no ties. When multiple entities have the same classification score, PerfMeas breaks ties based on their input order, and therefore different runs of PerfMeas with different input orders of the entities can produce different AUPRC values (i.e., Issue ➃). The PRC visualization also has these issues (i.e., Issue I).

#### PRROC

PRROC [17] is an R package specializing in constructing the ROC and PRC and computing the areas under them (Supplementary Table 1). PRROC provides two options (Table 1). The first option uses straight lines to connect adjacent anchor points when there are no ties and the discrete expectation method for handling ties (Supplementary Figures 4f, 5f, 6f, and 7f). The second option uses continuous expectation method no matter whether there are ties or not (Supplementary Figures 4g, 5g, 6g, and 7g). For both options, we used the pr.curve() function to construct the PRC and compute AUPRC, although with different parameter values to specify the options, and the plot() function to visualize the PRC.

PRROC constructs PRC by defining a finite number of points on it. When there are more than 100 actual positives, the PRCs constructed by the two methods are identical, both based on Formula 1 with *tp* set to 1. Otherwise, the continuous expectation method of PRROC places points at regular intervals along the x-axis, with an increase of 0.01 recall between every two adjacent points. When using the continuous expectation method, PRROC computes the AUPRC directly using Formula 3 rather than computing the area under the PRC it constructs.

We discovered a programming bug (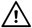) in the visualization of PRC when the discrete expectation method is used to handle ties. The bug is due to the inexact nature of floating-point number computation, which creates errors in sorting (Supplementary Figure 20). The part of the code that contains this bug is not used in the calculation of AUPRC, and therefore the correctness of AUPRC is not affected by it.

#### TensorFlow

TensorFlow [21] is a machine learning platform in Python and C++, most commonly used for deep learning (Supplementary Table 1). We used the tensorflow.keras.metrics.AUC() function to compute AUPRC. It computes AUPRC using the continuous expectation method without providing PRC visualizations (Table 1). We conceptually derive the AUPRC that corresponds to the AUPRC computed (Supplementary Figures 4h, 5h, 6h, and 7h).

We give three cautionary notes (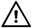) about using TensorFlow to compute AURPC. First, unlike most other tools, which use each unique classification score as a threshold, TensorFlow by default uses a user parameter (default to 200) to define the number of thresholds, which are then distributed evenly between zero and one. This uniform sampling approach is most accurate when the classification scores are distributed uniformly between zero and one, and therefore it is not suitable when most classification scores are concentrated in a small range of values. In addition, if the user parameter is set to a small value, the resulting AUPRC can deviate substantially from one computed by using each unique classification score as a threshold. Alternatively, TensorFlow provides an option for the user to provide a list of thresholds to use.

When different numbers of thresholds are used or when different threshold lists are provided, the resulting AUPRC computed can be different. We illustrate this issue in Supplementary Figure 21 based on an example provided on the manual page of TensorFlow that explains how AUPRC can be computed (https://tensorflow.google.cn/api_docs/python/tf/keras/metrics/AUC).

Second, due to the first cautionary note above, TensorFlow expects all classification scores to be between zero and one. Classification scores outside this range will not create a run time error, but they can make the AUPRC calculated inaccurate.

Third, instead of considering all entities with a classification score larger than or equal to the threshold as positive, which is what we described in the main text and implemented by most tools, TensorFlow considers only entities with a classification score larger than the threshold as positive (the smallest threshold that TensorFlow defines is a small negative value, such that entities with a zero classification score are all included in the last anchor point). We could not find any warnings about this cautionary notes on the official web site of TensorFlow.

#### precrec

precrec [20] is an R package specializing in constructing the ROC and PRC and computing the areas under them (Supplementary Table 1). We used the evalmod() function to compute AUPRC.

precrec uses the discrete expectation method to compute AUPRC (Table 1, Supplementary Figures 4i, 5i, 6i, and 7i). When there are no ties in classification scores, it creates points on the PRC in regular intervals of recall, where the length of each interval is a user parameter (default to 0.001), and computes the precision of these points using the expectation formula. When there are *x* entities with the same classification score, it first interpolates TP and FP to generate *x* points in total, converts them into recall and precision, and then creates points on the PRC in regular intervals of recall in the same way as it does when there are no ties.

#### TorchEval

TorchEval [23] is a library of the PyTorch machine learning framework, most commonly used in deep learning, for evaluating model performance (Supplementary Table 1). We used the tourcheval.metrics.BinaryAUPRC class to compute AUPRC. It uses the AP method to compute AUPRC without providing PRC visualizations (Table 1). We conceptually derive the AUPRC that corresponds to the AUPRC computed (Supplementary Figures 4j, 5j, 6j, and 7j).

We did not find any issues with the AUPRC calculation of TorchEval.

#### MLeval

MLeval [18] is an R package for evaluating machine learning models (Supplementary Table 1). We used the evalm() function to construct PRC and compute AUPRC. It uses straight lines to connect adjacent anchor points when there are no ties and the linear interpolation method for handling ties (i.e., Issue ➀) (Table 1, Supplementary Figures 4k, 5k, 6k, and 7k). It only connects anchor points, which means the PRC does not cover the area with a recall values smaller than that of the first anchor point (i.e., (i.e., Issue ➂. The PRC visualization has the same issues (i.e., Issue I)

#### yardstick

yardstick [19] is an R package that provides various metrics for quantifying how well a model fits a data set (Supplementary Table 1). It provides two options for computing AUPRC (Table 1), by the direct straight line/linear interpolation method (Supplementary Figures 4l, 5l, 6l, and 7l) and the AP method (Supplementary Figures 4m, 5m, 6m, and 7m), respectively. For the first option, we used the pr_curve() function to construct PRC and the pr_auc() function to compute AUPRC. For the second option, we used the average_precision() function to compute AUPRC.

For the first option, in addition to the issue of linear interpolation (i.e., Issue ➀), yardstick also always starts the PRC at (0, 1) (i.e., Issue ➁). The PRC visualization has the same issues (i.e., Issue I).

### The ROC and AUROC

Although this study focuses on issues related to PRC and AUPRC, all the tools for producing PRC and AUPRC we analyzed can also compute AUROC. We also found some tools that can compute AUROC but not AUPRC (Supplementary Table 1). Since ROC and PRC are conceptually highly related to each other and practically both commonly used, here we also explain the concepts behind ROC and AUROC and their implementations.

### Definition of the ROC and AUROC

The ROC plots the true positive rate (TPR) against the false positive rate (FPR) (Figure 1). In the most common way to produce the ROC, which is used by all the 12 tools we analyzed (Supplementary Figure 1), each unique classification score is used as a threshold to define an anchor point of the ROC. When no two entities have the same classification score, two adjacent anchor points are connected directly by a straight line. The starting point of the ROC is obvious from its definition: if a threshold larger than the classification scores of all the entities is used, none of the entities is predicted to belong to the target class. In that situation, both the TPR and the FPR are zero. Therefore, the ROC always starts at (0, 0). When multiple entities have the same classification score, they together define a single anchor point. This anchor point is connected to the previous anchor point by a direct straight line, assuming that all possible orders of these entities are equally likely. In other words, ties are handled by taking the expectation [45] (Supplementary Figure 22). To understand why the expectation curve is linear in the case of ROC but non-linear in the case of PRC, we note that in the definitions of TPR and FPR, the denominator is a constant (total number of actual positives and total number of actual negatives, respectively) and therefore TPR is directly proportional to the number of true positives so far and FPR is directly proportional to the number of false positives so far. In contrast, in the definition of precision, the denominator is the total number of true positives and false positives so far. Therefore, precision is not directly proportional to the number of true positives.

All the tools we surveyed handled ties in this way. The only difference between them is whether additional points are interpolated between the anchor points. These additional points do not change the shape of the ROC or the calculation of AUROC, but they can be useful for computing confidence intervals [20].

A recent study argued that when there are ties, directly connecting adjacent anchor points by straight lines is not appropriate when the classifier produces discrete predictions [46]. This is still under debate.

### Algorithms for plotting the ROC

There are two commonly used algorithms for plotting the ROC, which produce identical results but differ from each other slightly in terms of their implementations.

The first algorithm considers the segment of the ROC directly caused by each entity one by one when there are no ties. Specifically, all entities are first sorted in descending order of their classification scores. Then each entity on the sorted list is visited sequentially. If it is a true positive, the ROC goes up vertically by 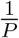, where *P* is the total number of actual positives among all the entities. If it is a false positive, the ROC goes right horizontally by 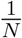, where *N* is the total number of actual negatives among all the entities. When multiple entities have the same classification score, the previous anchor point is connected to a new anchor point that has an increase of 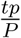 TPR and an increase of 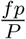 FPR, where *tp* and *fp* are the number of actual positives and actual negatives among these entities (Supplementary Figure 23a, b).

The second algorithm uses the definition of ROC directly. It computes the coordinates of each anchor point based on the TPR and FPR when its threshold is used to predict entities that belong to the positive class (Supplementary Figure 23a, c). To determine the anchor points, most of the tools also sort the entities in descending order of their classification scores.

### Algorithms for computing AUROC

There are two common algorithms for computing AUROC, which produce identical results [47, 48].

The first one accumulates the change of AUROC by summing up the areas of all the trapezoids (Supplementary Figure 24). Each accumulation happens either when an anchor point has a FPR larger than the previous anchor point (the vertical trapezoidal rule, Supplementary Figure 24a) or when an anchor point has a TPR larger than the previous anchor point (the horizontal trapezoidal rule, Supplementary Figure 24b). Most of the tools we surveyed use the vertical trapezoidal rule, including ROCR, pROC [49], scikit-learn, PRROC, TensorFlow, plotROC [50], precrec (default mode), TorchEval, MLeval, and yardstick (Supplementary Table 3). Weka uses the horizontal trapezoidal rule and further divides each trapezoid into a triangle and a rectangle.

The second method computes the Wilcoxon-Mann-Whitney statistic and then converts it to AUROC (Supplementary Figure 25, to be explained in the next section below). Precrec (aucroc mode) and PerfMeas compute AUROC in this way with slightly different procedures. There are some other metrics that are either closely related to the Wilcoxon-Mann-Whitney statistic or identical to it under certain circumstances. These metrics can also be used to compute AUROC. For example, the concordance index, which is an evaluation metric frequently used in survival analyses, is identical to the Wilcoxon-Mann-Whitney statistic when predictions are binary [51].

#### Relationship between the Wilcoxon-Mann-Whitney statistic and AUROC

Suppose there are two lists of real values. The Wilcoxon-Mann-Whitney statistic quantifies the difference between them based on the sum of the ranks of the values from each list when the two lists are merged. Conceptually, if the two lists have comparable values, their rank sums should be similar. Using this statistic, the Wilcoxon-Mann-Whitney test (also called “rank-sum test” and “U test”) compares two distributions based on a sample from each of them (i.e., the two lists), under the null assumption that a value randomly sampled from the first distribution has equal chance to be larger than or smaller than a value randomly sampled from the second distribution. By considering only the ranks but not the original values, the Wilcoxon-Mann-Whitney test is not specific to any parametric family of distributions.

Mathematically, the Wilcoxon-Mann-Whitney statistic based on the values on the first list is defined as

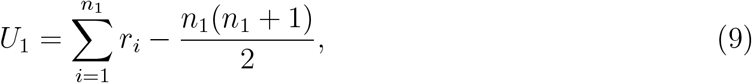

where *n*_1_ is the number of values on the first list and *r*_*i*_ is the rank of its *i*-th largest value in the merged list (largest value in the merged list has rank 1). Notice that if the merged list contains duplicated values, they all share the same average rank. For example, if the four largest values on the merged list are the same, they all have a rank of 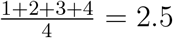.

According to the definition, the Wilcoxon-Mann-Whitney statistic is the difference between two components. The first component is the rank sum of the values in the merged list. The second component is the smallest possible rank sum of *n*_1_ values in the merged list, which happens when all values on the first list are larger than all values on the second list. The Wilcoxon-Mann-Whitney statistic therefore quantifies how far the actual rank sum is from the minimum.

To see how *U*_1_ is related to AUROC, we treat the first list as the classification scores of the actual positive entities and the second list as the classification scores of the actual negative entities. Therefore, *n*_1_ = *P*. For simplicity of notations and without loss of generality, we assume that both lists are sorted in descending order.

First, we consider the situation that all entities have unique classification scores. The contribution of the *i*-th entity on the actual positive list to the AUROC is the area of a rectangular stripe with a height of one unit and a width of *N* − *f*_*i*_ units, where *f*_*i*_ is the number of actual negatives having a higher classification score than the *i*-th actual positive entity (Supplementary Figure 25a). After normalizing by the total number of actual positives and total number of actual negatives, the area of this rectangle is:

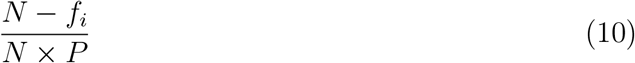

Summing over the contributions of all the actual positive entities, we get the AUROC:

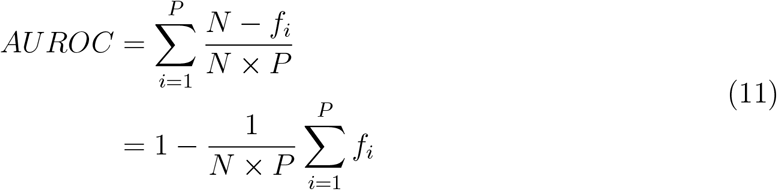

Now, the rank of an actual positive entity in the merged list is equal to the number of entities with a classification score higher than or equal to it, including itself, other actual positive entities, and actual negative entities. Therefore, *r*_*i*_ = *i* + *f*_*i*_. This means the formula of AUROC can be re-written as follows:

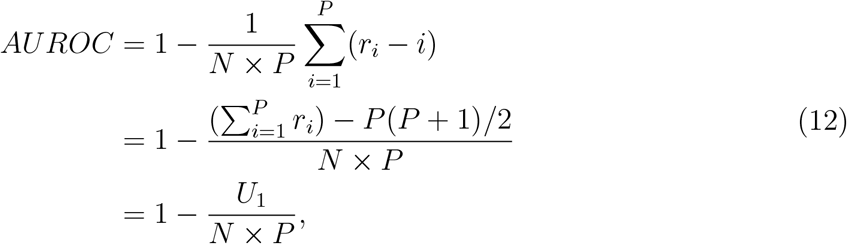

which provides a way to compute AUROC from the Wilcoxon-Mann-Whitney statistic [52]. Next, we consider the situation that some entities have the same classification score. Suppose there is a group of entities that have the same classification score, among which there are *p* actual positives and *n* actual negatives. Suppose also that in the list of actual positives and the merged list of actual positives and actual negatives (both lists sorted in descending order of classification scores), the first entry with this classification score is the *i*-th entry and the *r*_*i*_-th entry, respectively. As mentioned above, each entity in this group has an average rank of 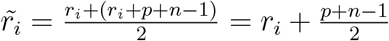 in the merged list. The total contribution of this group to the AUROC is the area of a trapezoid with a height of *p* units, a width of *N* − *f*_*i*_ on the lower, longer side, and a width of *N* − *f*_*i*_ − *n* on the upper, shorter side (Supplementary Figure 25b). After normalizing by the total number of actual positives and total number of actual negatives, the area of this trapezoid is:

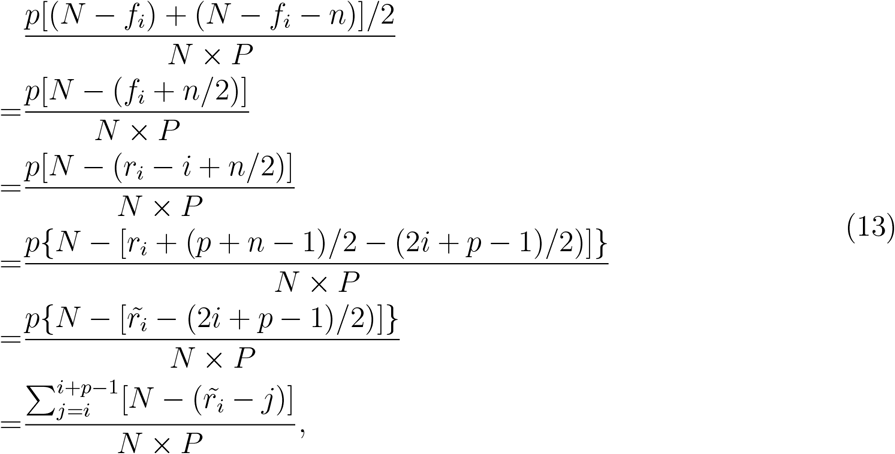

which has the same form as the summation formula for computing the AUROC based on the contributions of individual actual positive entities (Formula 11). Therefore, AUROC can still be computed from the Wilcoxon-Mann-Whitney statistic using Formula 12 when all entities in ties are given their average rank.

#### Special notes on the implementations of AUROC calculation by different tools

**Precrec** (aucroc mode) computes AUROC from the Wilcoxon-Mann-Whitney statistic essentially following our description above, except that it sorts classification scores in ascending order and adjusts the formulas accordingly.

**PerfMeas** also computes AUROC from the Wilcoxon-Mann-Whitney statistic essentially following our description above, except that it sums over contributions of individual actual negatives and adjusts the formulas accordingly.

**TensorFlow**’s AUROC calculation also uses the designs that we discussed in the three cautionary notes for its AUPRC calculation. Therefore, the accuracy of the AUROC computed also depends on the distribution of classification scores. In the same official example provided by TensorFlow (Supplementary Figure 21), the correct AUROC computed by having 3 thresholds seems only a coincidence.

## Supplementary tables

**Supplementary Table 1:**
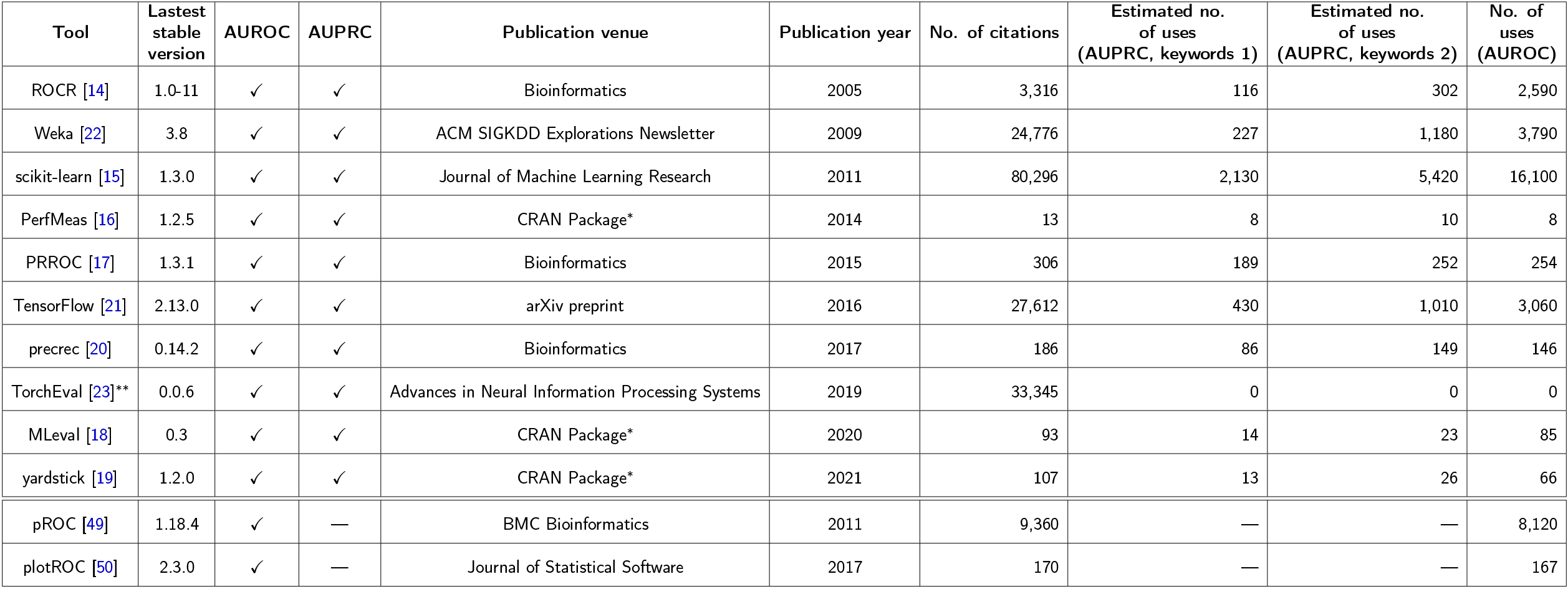
List of tools evaluated in this study. The 10 tools that can compute AUPRC are listed first, followed by 2 tools that can only compute AUROC but not AUPRC. In each category, the tools are ordered by year of corresponding publications. ^*^For the CRAN packages, there were no associated formal papers available, leading to some publications where these packages were used without formal citation, making it difficult for Google Scholar to accurately count their citations. To resolve this problem, we conducted a manual search for each package name on Google Scholar and verified whether the resulting papers cited these packages. ^**^TorchEval has been embedded into PyTorch officially (https://pytorch.org/torcheval/stable/) without a formal publication; we collected the number of citations of the original publication of PyTorch instead. As there is no stable version of TorchEval yet, we collected its latest version 0.0.6.

**Supplementary Table 2:** Consequences of the five issues in computing AUPRC.

| Issue | Can lead to overly-optimistic AUPRC |  | Can flip the order of two AUPRCs |  |
| --- | --- | --- | --- | --- |
|  | With ties | Without ties | With ties | Without ties |
| ① | Yes | No | Yes | No |
| ② | Yes | No | Yes | No |
| ③ | No | No | Yes | Yes |
| ④ | Yes | No | Yes | No |
| ⑤ | Yes | Yes | Yes | Yes |

**Supplementary Table 3:**
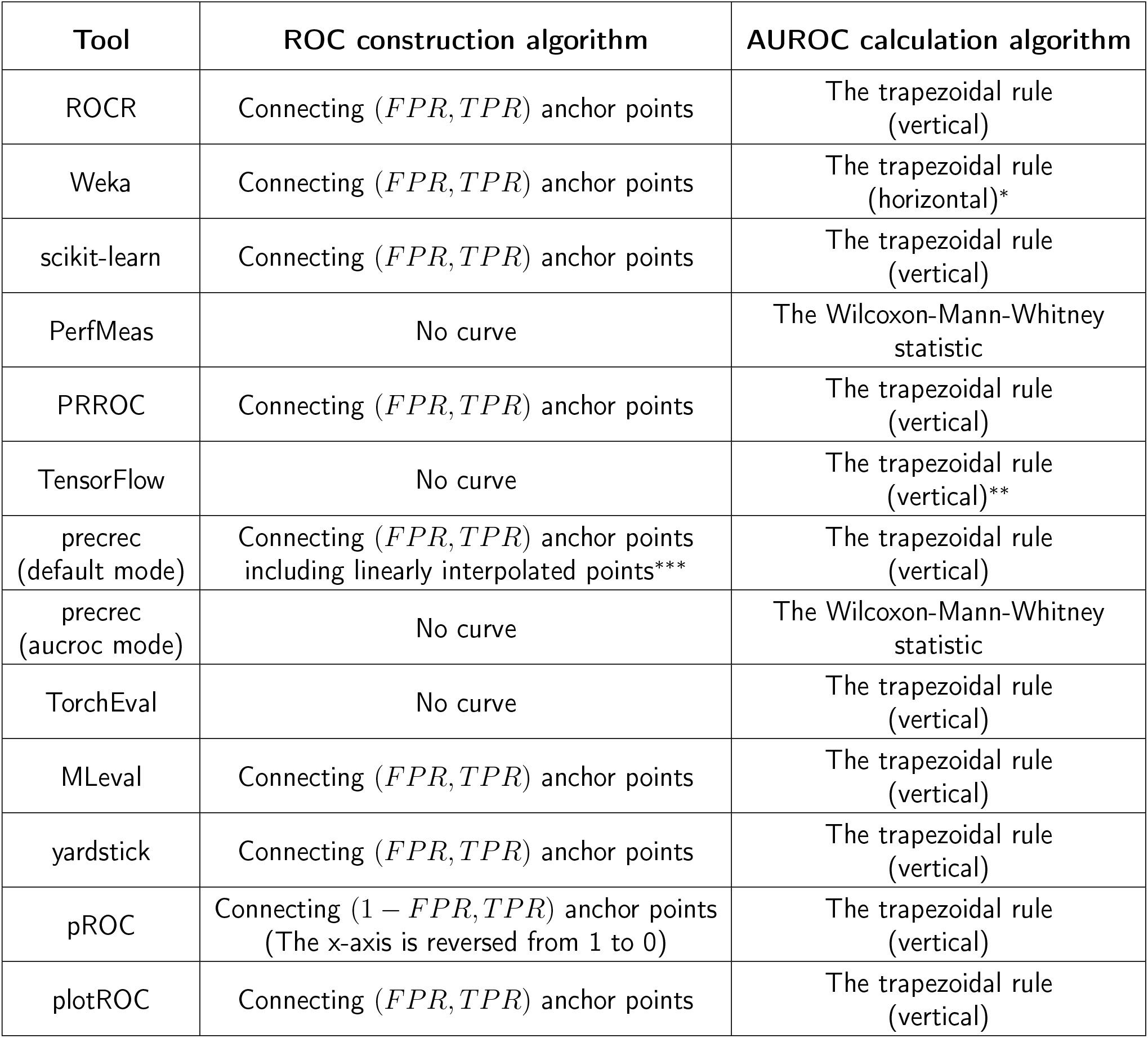
Algorithms used by the 12 tools we surveyed to construct ROC and compute AUROC. ^*^Weka states that it uses the Wilcoxon-Mann-Whitney statistic to compute AUROC in its source code. However, by checking the logic implemented by the source code and its naming scheme (i.e., the variable name is “area”), we believe it actually uses the trapezoidal rule (horizontal). ^**^TensorFlow specifies the Riemann summation method in its source code when it computes AUROC, which is equivalent to the trapezoidal rule (vertical). ^***^Precrec employs linear interpolation with the interpolated points explicitly created to construct the ROC.

## Supplementary figures

**Supplementary Figure 1:**
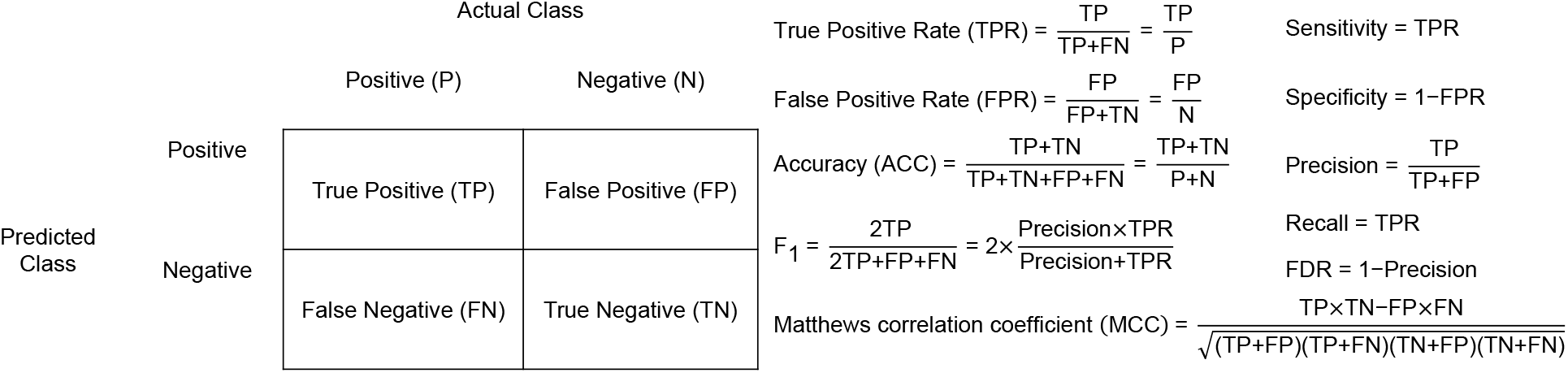
The confusion matrix and definitions of some commonly used measures calculated from it. In these definitions, P, N, TP, FP, FN, and TN are the numbers of entities in these categories based on a specific classification threshold.

**Supplementary Figure 2:**
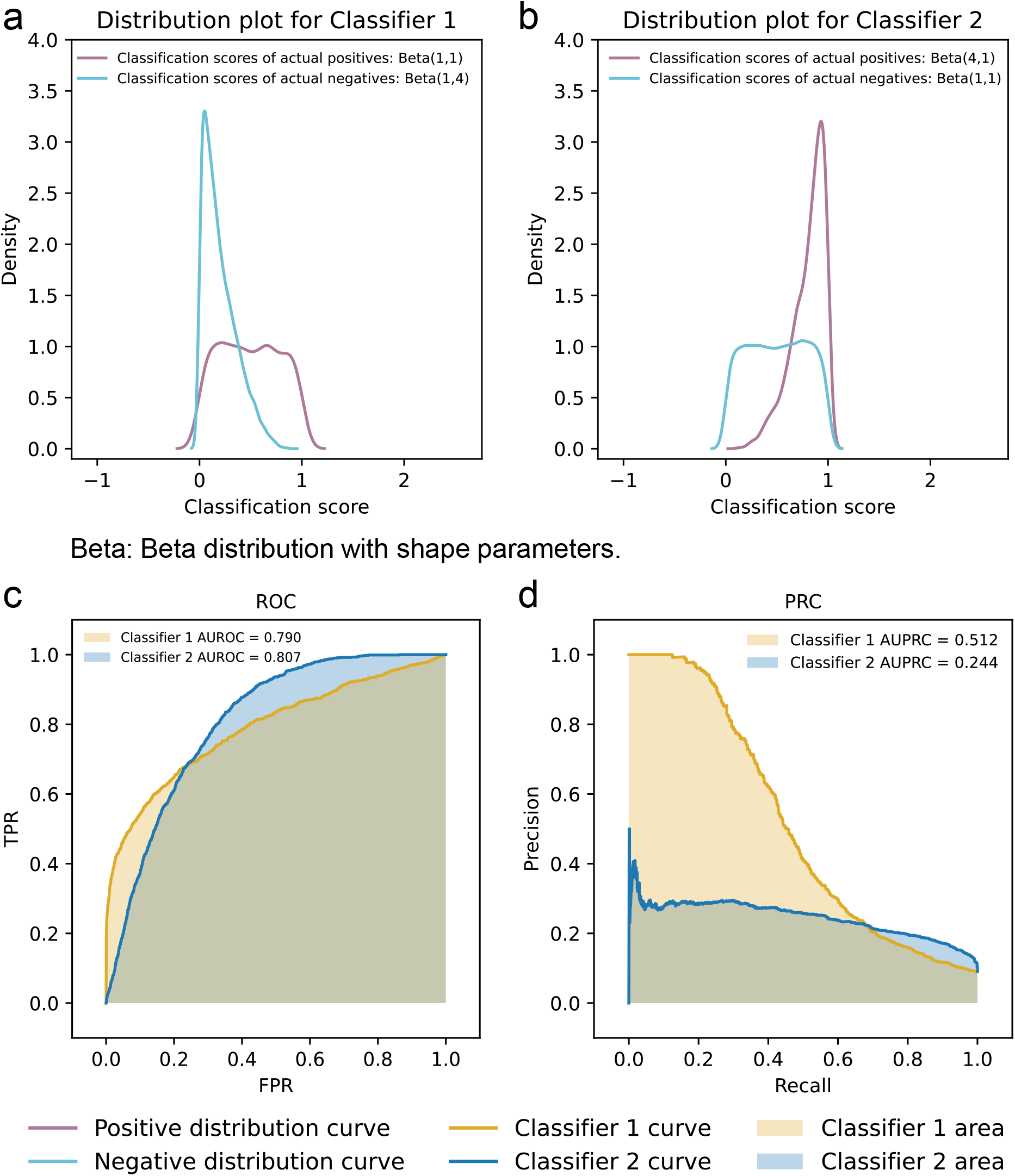
A demonstration that AUPRC is more sensitive to performance differences than AUROC when the target class has few entities and there are errors among the top predictions. **a**,**b** Classification scores produced by two classifiers simulated using Beta distributions [1]. A larger proportion of actual positives are predicted among the entities receiving the highest classification scores produced by Classifier 1 than Classifier 2. **c** The ROCs and AUROCs of the two classifiers. Classifier 1 has a slightly smaller AUROC than Classifier 2. **d** The PRCs and AUPRCs of the two classifiers. Classifier 1 has a much larger AUPRC than Classifier 2.

**Supplementary Figure 3:**
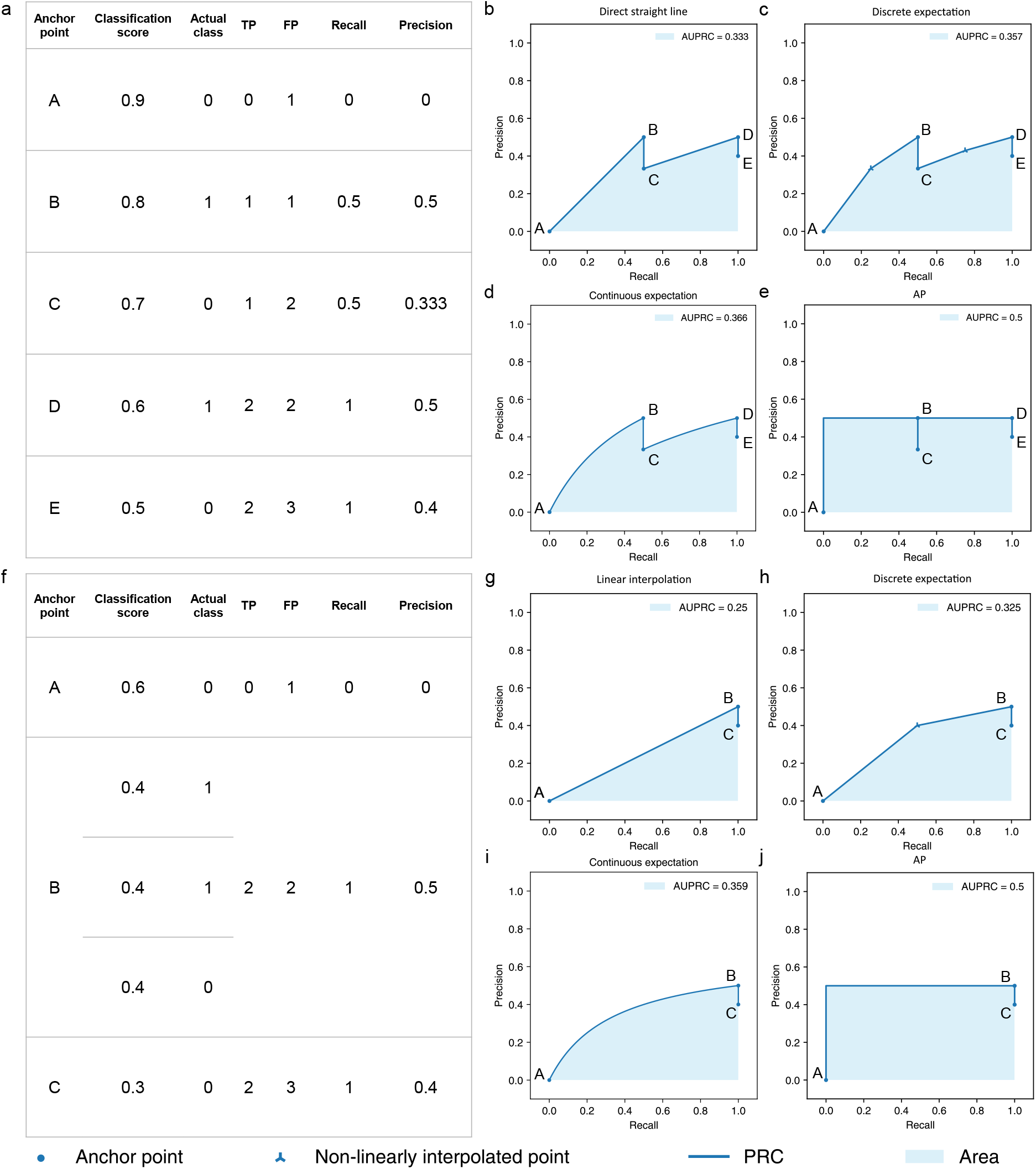
Extra examples that illustrative different methods for connecting adjacent anchor points on the PRC. **a** An illustrative data set with no two entities receiving the same classification score. **b-e** Different methods for connecting adjacent anchor points when there are no ties in classification scores, namely **b** direct straight line, **c** discrete expectation, **d** continuous expectation, and **e** AP. **f** An illustrative data set with different entities receiving the same classification score. Each group of entities with the same classification score defines a single anchor point (A, B, and C). **g-j** Different methods for connecting anchor point B to its previous anchor point, A, namely **g** linear interpolation, **h** discrete expectation, **i** continuous expectation, and **j** AP. In **c** and **h**, *tp* is set to 0.5 and 1 in Formula 1, respectively (Supplementary text).

**Supplementary Figure 4:**
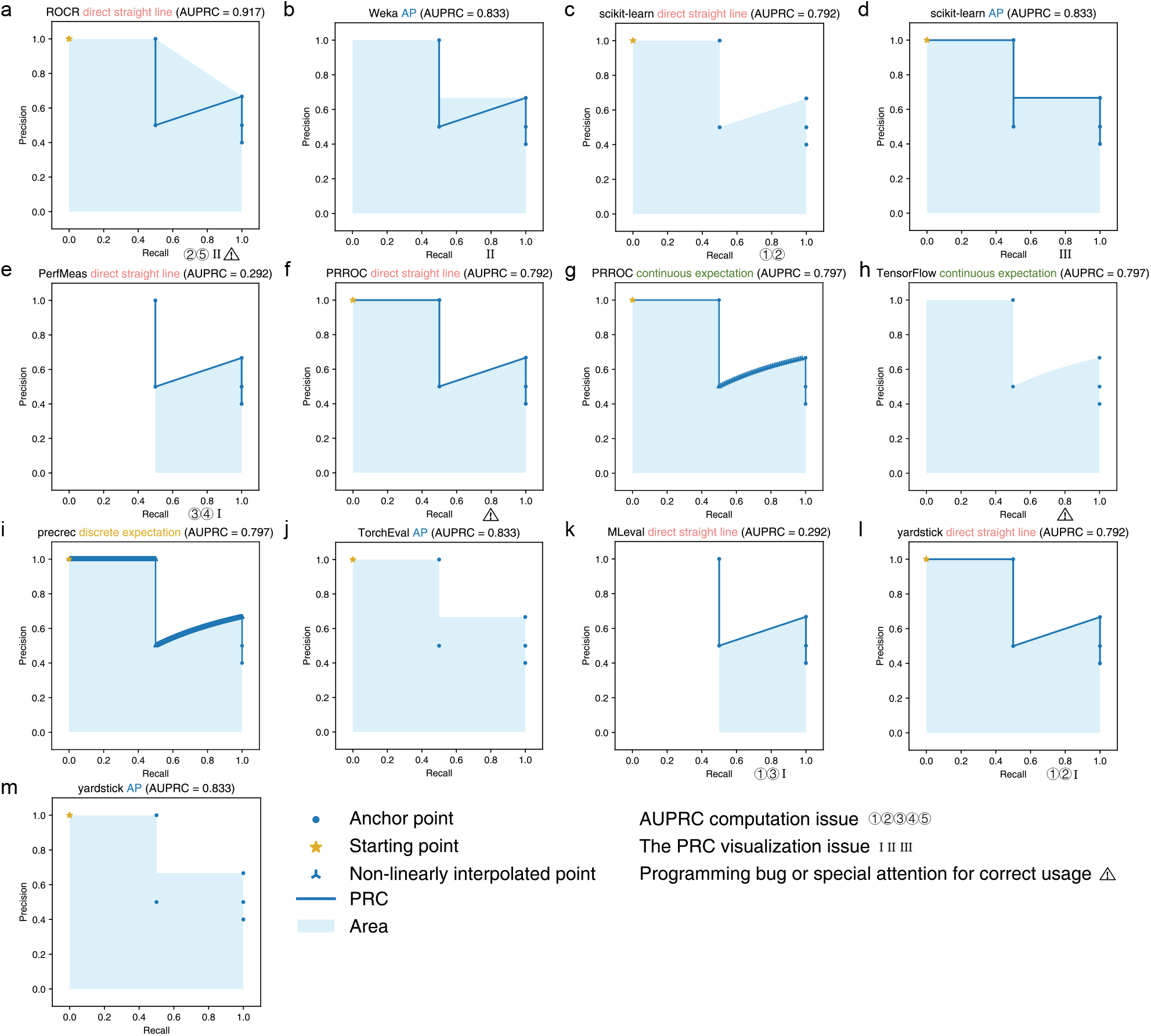
The PRCs and AUPRCs of the illustrative example in Figure 1a produced by the 10 tools. **a** ROCR (direct straight line). **b** Weka (AP). **c** scikit-learn (direct straight line). **d** scikit-learn (AP). **e** PerfMeas (direct straight line). **f** PRROC (direct straight line). **g** PRROC (continuous expectation). **h** TensorFlow (continuous expectation). **i** precrec (discrete expectation). **j** TorchEval (AP). **k** MLeval (direct straight line). **l** yardstick (direct straight line). **m** yardstick (AP).

**Supplementary Figure 5:**
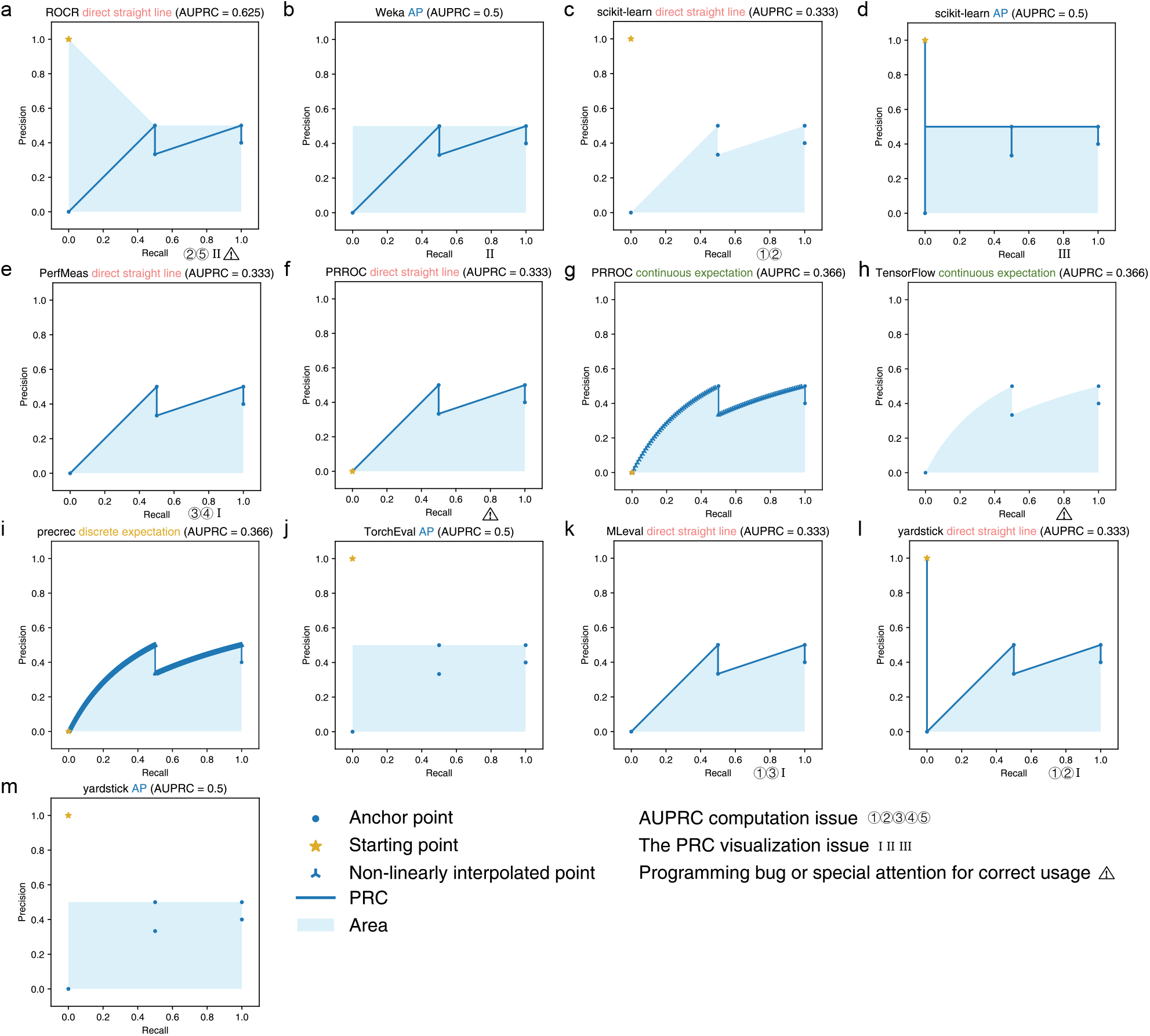
The PRCs and AUPRCs of the illustrative example in Supplementary Figure 3a produced by the 10 tools. **a** ROCR (direct straight line). **b** Weka (AP). **c** scikit-learn (direct straight line). **d** scikit-learn (AP). **e** PerfMeas (direct straight line). **f** PRROC (direct straight line). **g** PRROC (continuous expectation). **h** TensorFlow (continuous expectation). **i** precrec (discrete expectation). **j** TorchEval (AP). **k** MLeval (direct straight line). **l** yardstick (direct straight line). **m** yardstick (AP).

**Supplementary Figure 6:**
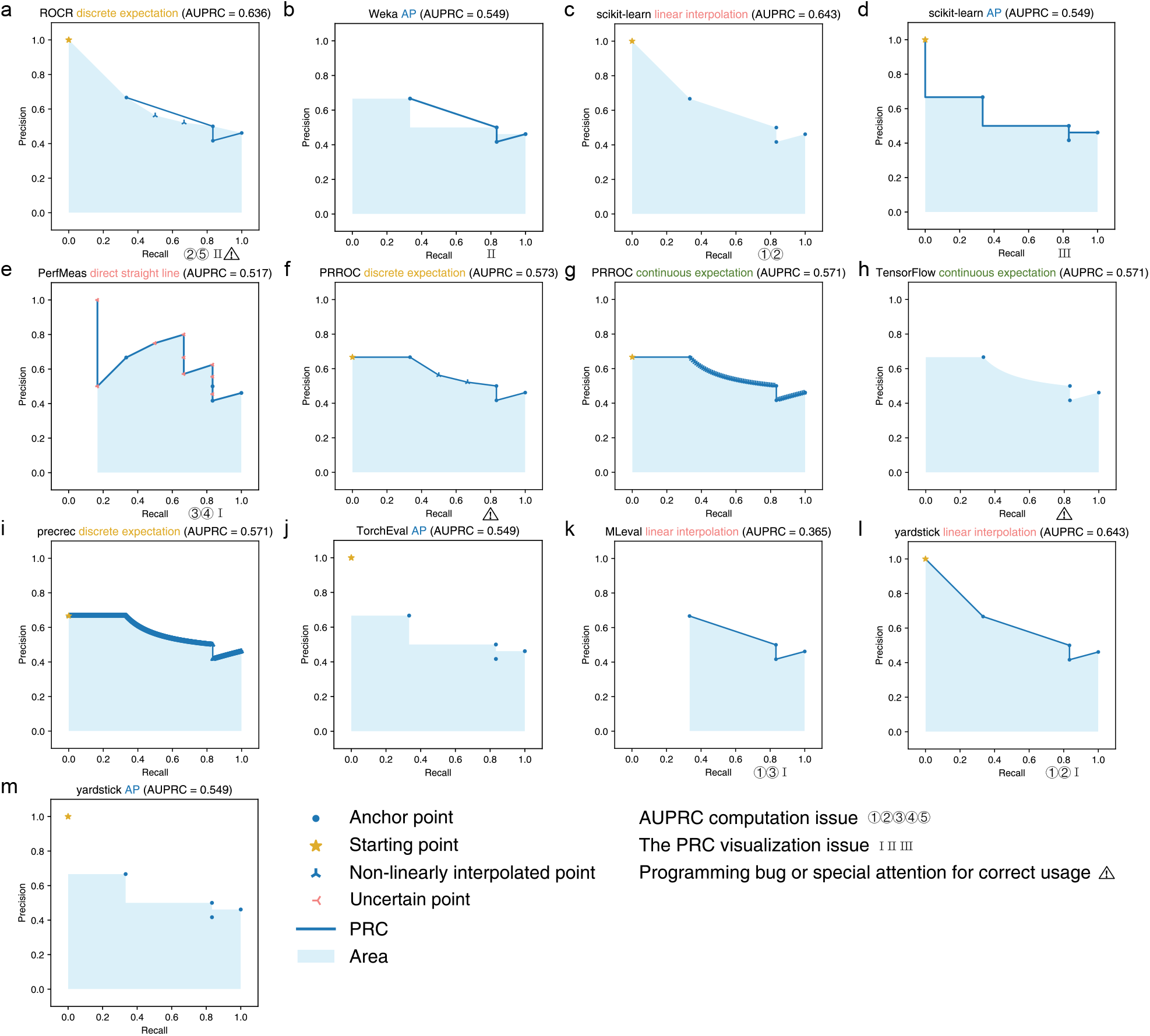
The PRCs and AUPRCs of the illustrative example in Figure 1f produced by the 10 tools. **a** ROCR (discrete expectation). **b** Weka (AP). **c** scikit-learn (linear interpolation). **d** scikit-learn (AP). **e** PerfMeas (direct straight line). **f** PRROC (discrete expectation). **g** PRROC (continuous expectation). **h** TensorFlow (continuous expectation). **i** precrec (discrete expectation). **j** TorchEval (AP). **k** MLeval (linear interpolation). **l** yardstick (linear interpolation). **m** yardstick (AP). The uncertain points on the PRC produced by PerfMeas are caused by its dependency on the input order of entities in ties.

**Supplementary Figure 7:**
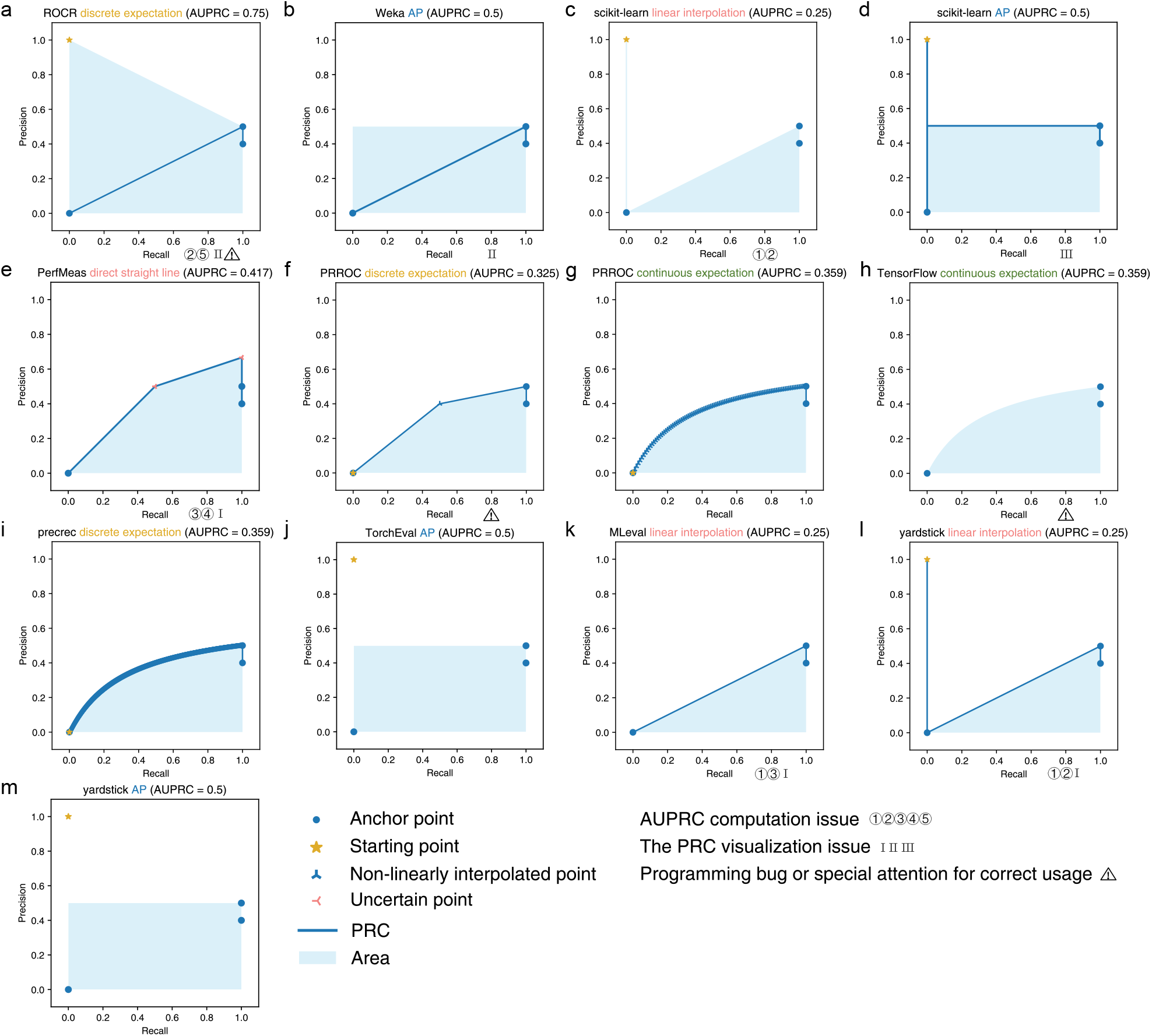
The PRCs and AUPRCs of the illustrative example in Supplementary Figure 3f produced by the 10 tools. **a** ROCR (discrete expectation). **b** Weka (AP). **c** scikit-learn (linear interpolation). **d** scikit-learn (AP). **e** PerfMeas (direct straight line). **f** PRROC (discrete expectation). **g** PRROC (continuous expectation). **h** TensorFlow (continuous expectation). **i** precrec (discrete expectation). **j** TorchEval (AP). **k** MLeval (linear interpolation). **l** yardstick (linear interpolation). **m** yardstick (AP). The uncertain points on the PRC produced by PerfMeas are caused by its dependency on the input order of entities in ties.

**Supplementary Figure 8:**
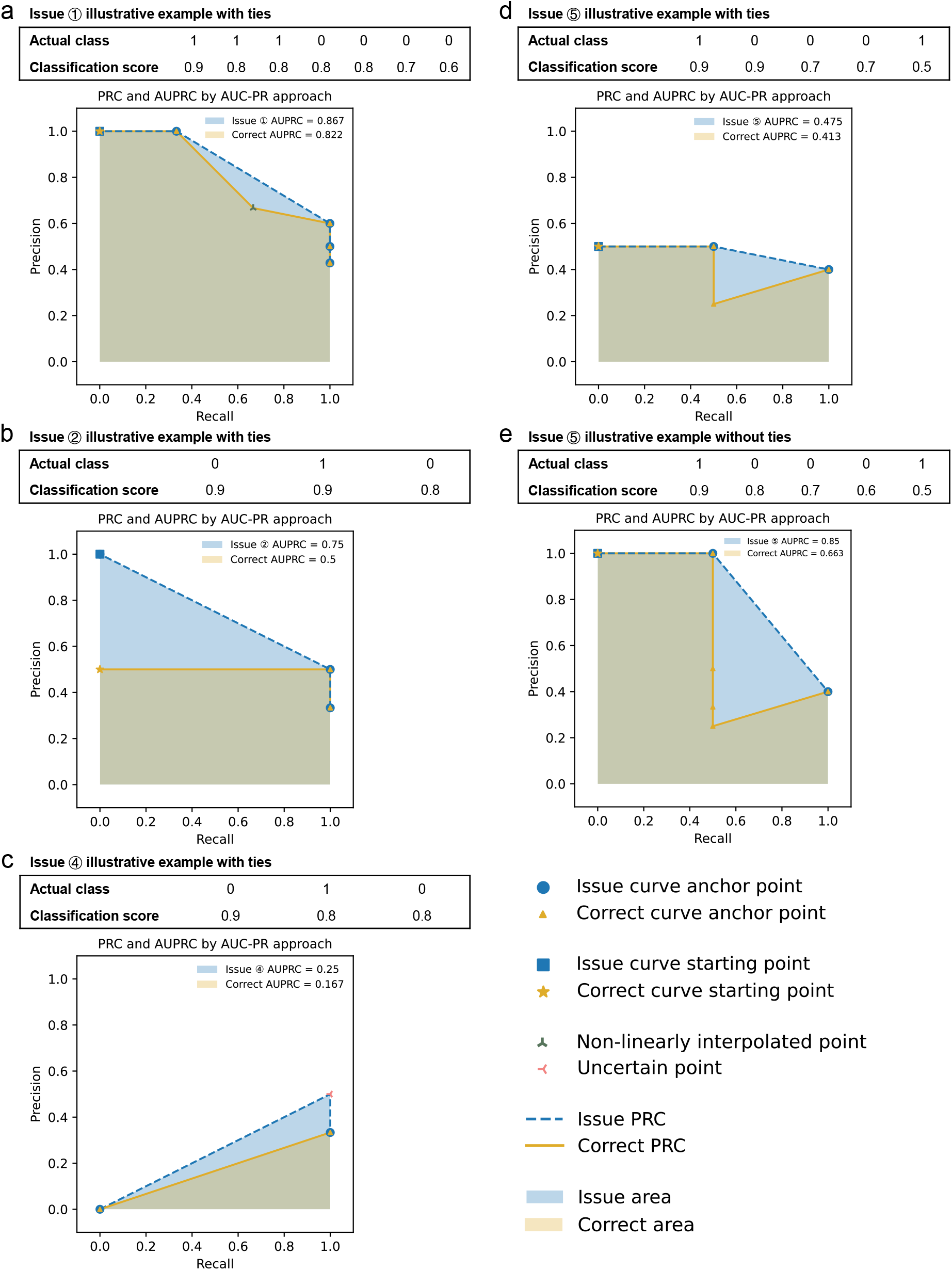
Illustrative examples of overly optimistic AUPRC values caused by the issues. **a** Issue ➀. **b** Issue ➁. **c** Issue ➃. **d-e** Issue ➄ with (**d**) or without (**e**) ties in classification scores. The correct PRCs and AUPRCs were generated by the bug-fixed version of PRROC (option that uses the discrete expectation method to handle ties).

**Supplementary Figure 9:**
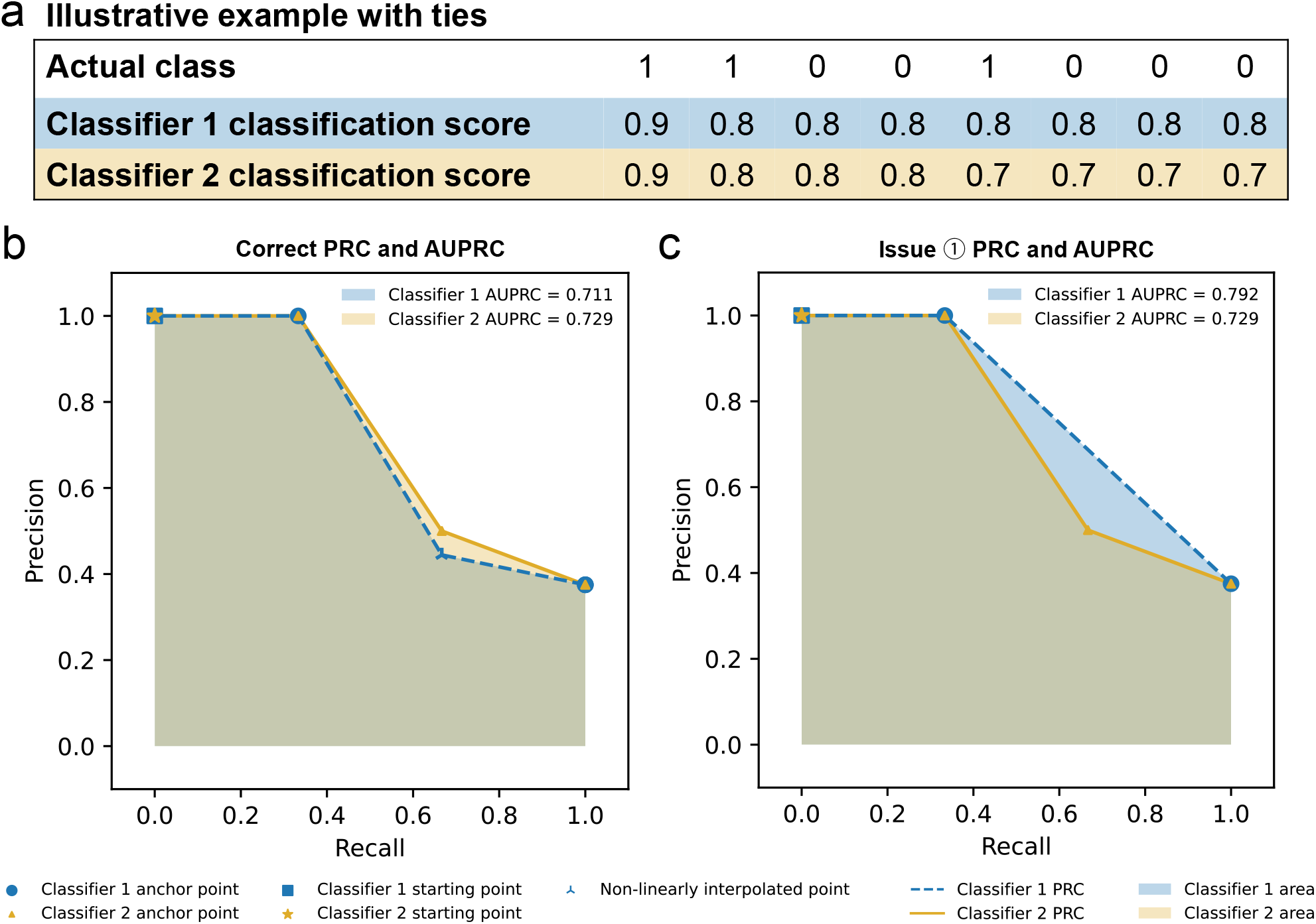
An illustrative example of flipping the orders of two classifiers based on their AUPRC values due to Issue ➀ when there are ties in classification scores. **a** An illustrative data set. **b** The PRCs and AUPRCs of two classifiers produced by the bug-fixed version of PRROC (option that uses the discrete expectation method to handle ties). **c** The PRCs and AUPRCs of the two classifiers with Issue ➀.

**Supplementary Figure 10:**
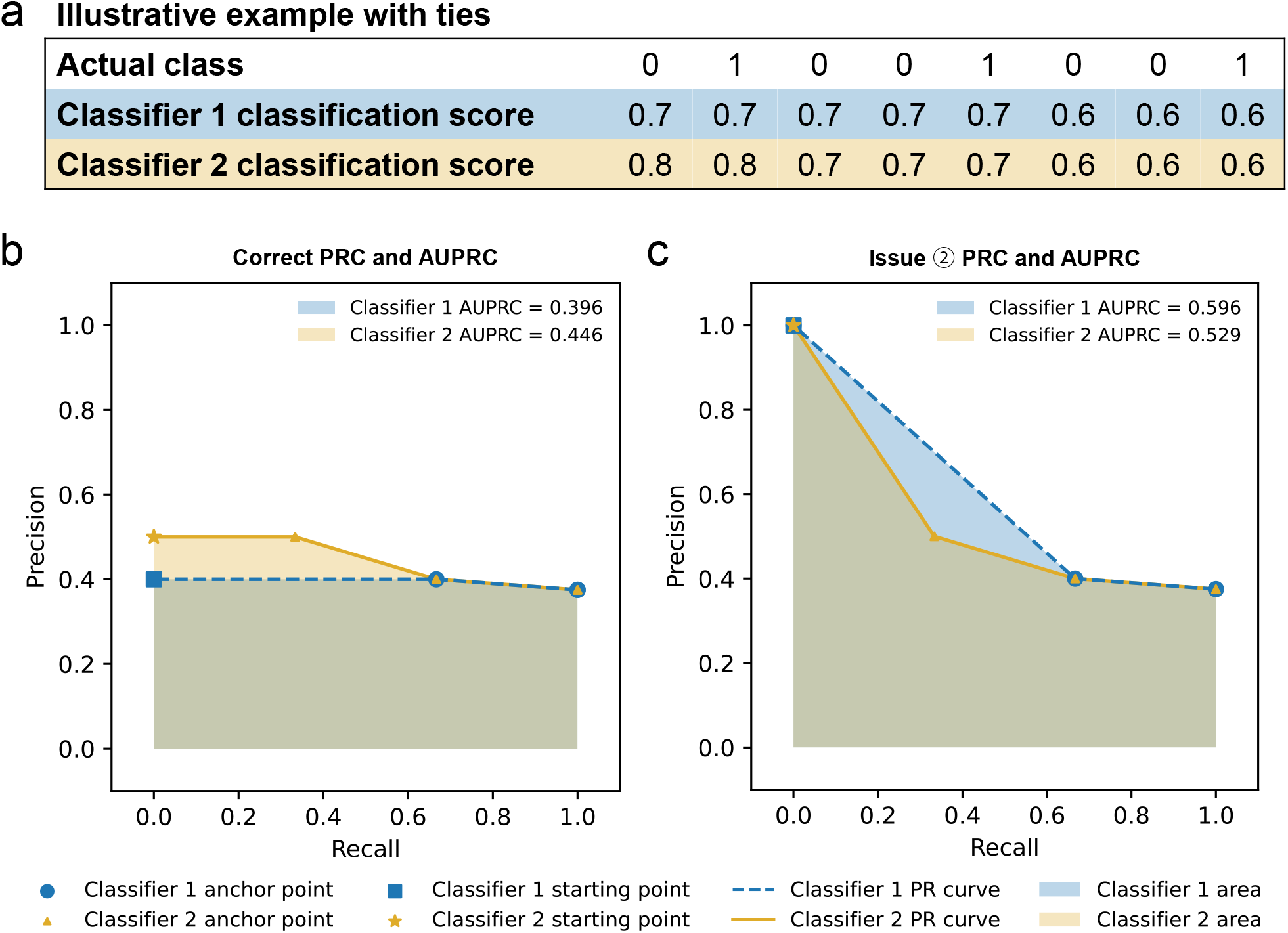
An illustrative example of flipping the orders of two classifiers based on their AUPRC values due to Issue ➁ when there are ties in classification scores. **a** An illustrative data set. **b** The PRCs and AUPRCs of two classifiers produced by the bug-fixed version of PRROC (option that uses the discrete expectation method to handle ties). **c** The PRCs and AUPRCs of the two classifiers with Issue ➁.

**Supplementary Figure 11:**
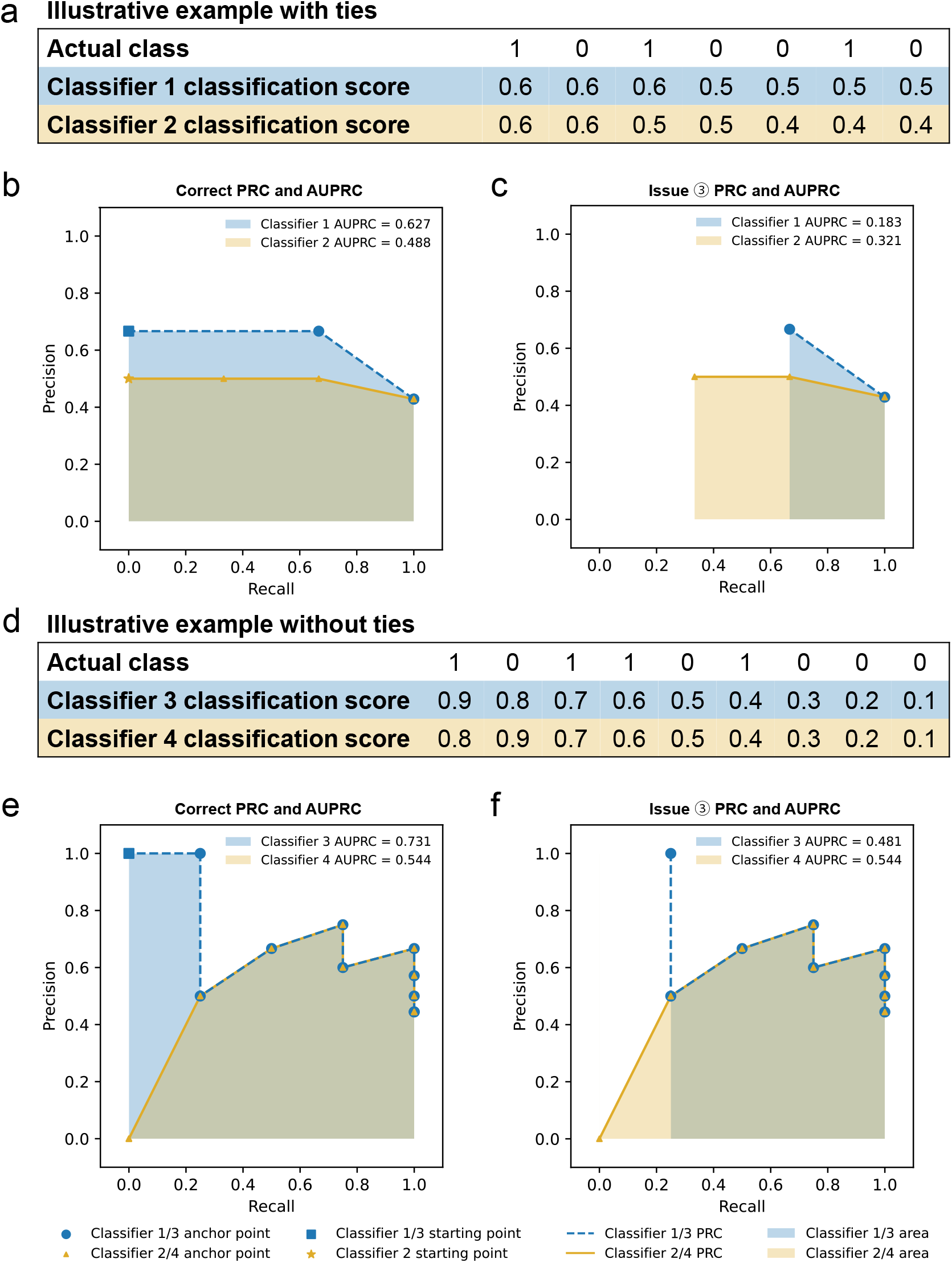
Illustrative examples of flipping the orders of two classifiers based on their AUPRC values due to Issue ➂ with and without ties in classification scores. **a**,**d** Illustrative data sets with (**a**) and without (**d**) ties. **b**,**e** The PRCs and AUPRCs of two classifiers produced by the bug-fixed version of PRROC (option that uses the discrete expectation method to handle ties) on the illustrative data set with (**b**) and without (**e**) ties. **c**,**f** The PRCs and AUPRCs of the two classifiers with Issue ➂ based on the illustrative data set with (**c**) and without (**f**) ties.

**Supplementary Figure 12:**
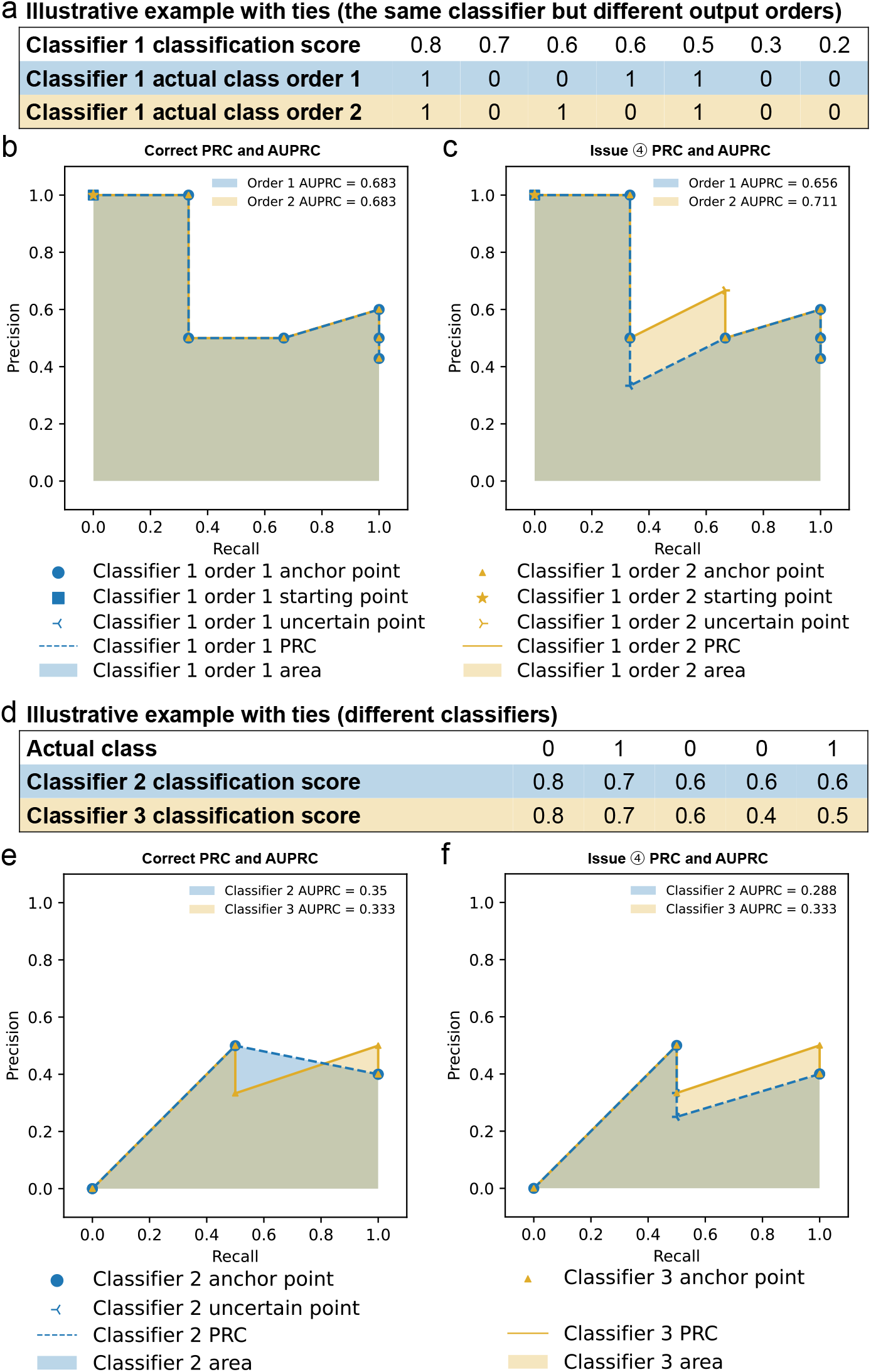
Illustrative examples of Issue ➃. **a** An illustrative data set with ties in classification scores produced by a classifier and two different input orders of the entities. **b** The correct PRC and AUPRC, which stays the same regardless of the input order of the entities. **c** The PRCs and AUPRCs produced if entities with the same classification score are ordered based on their input order. **d** An illustrative data set with ties in classification scores produced by one of the two classifiers. **e**,**f** The PRCs and AUPRCs of the two classifiers with Issue ➃.

**Supplementary Figure 13:**
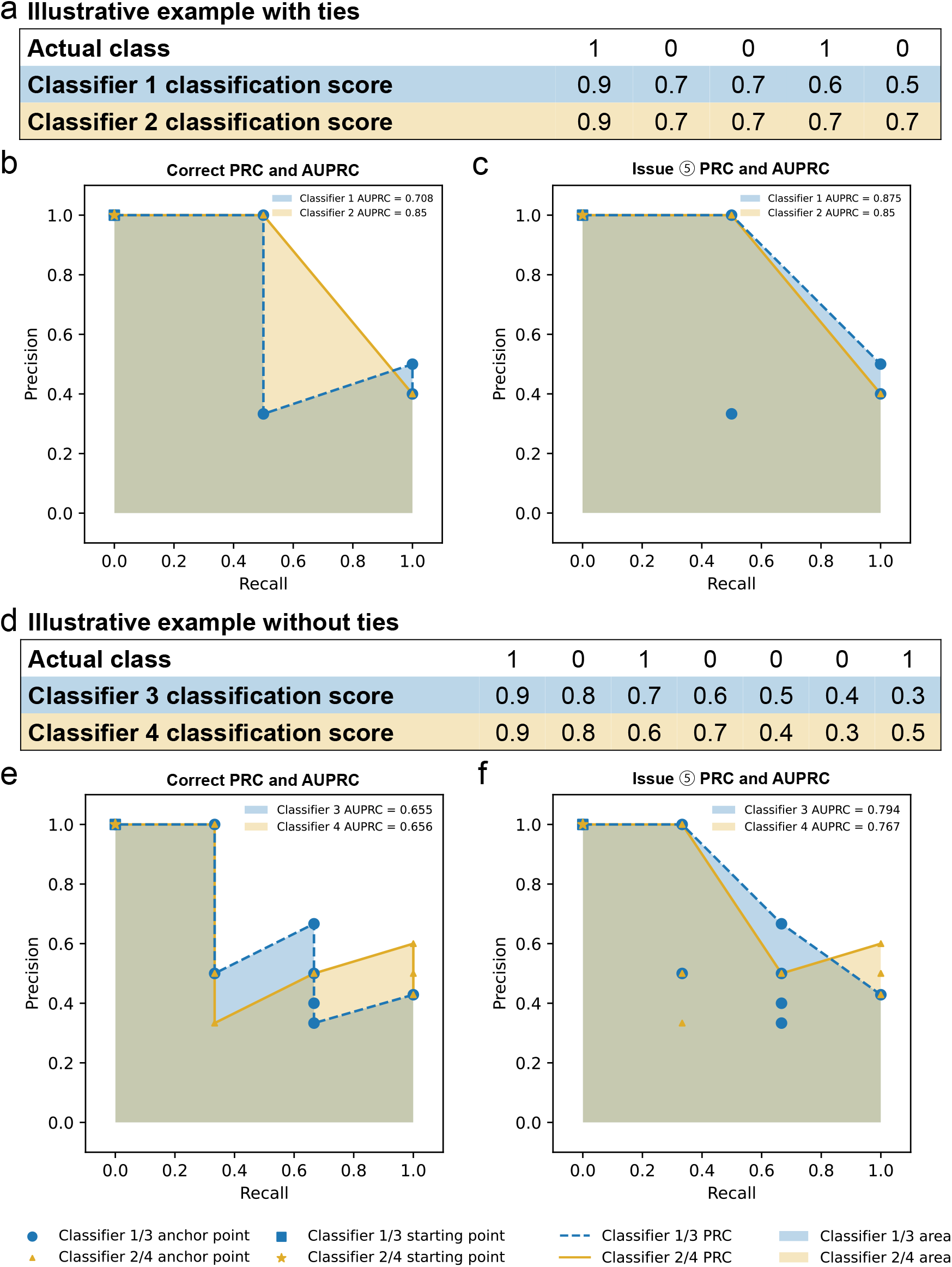
Illustrative examples of flipping the order of two classifiers based on their AUPRC values due to Issue ➄ with and without ties in classification scores. **a**,**d** Illustrative data sets with (**a**) and without (**d**) ties. **b**,**e** The PRCs and AUPRCs of two classifiers produced by the bug-fixed version of PRROC (option that uses the discrete expectation method to handle ties). **c**,**f** The PRCs and AUPRCs of the two classifiers with Issue ➄ on the illustrative data set with (**c**) and without (**f**) ties.

**Supplementary Figure 14:**
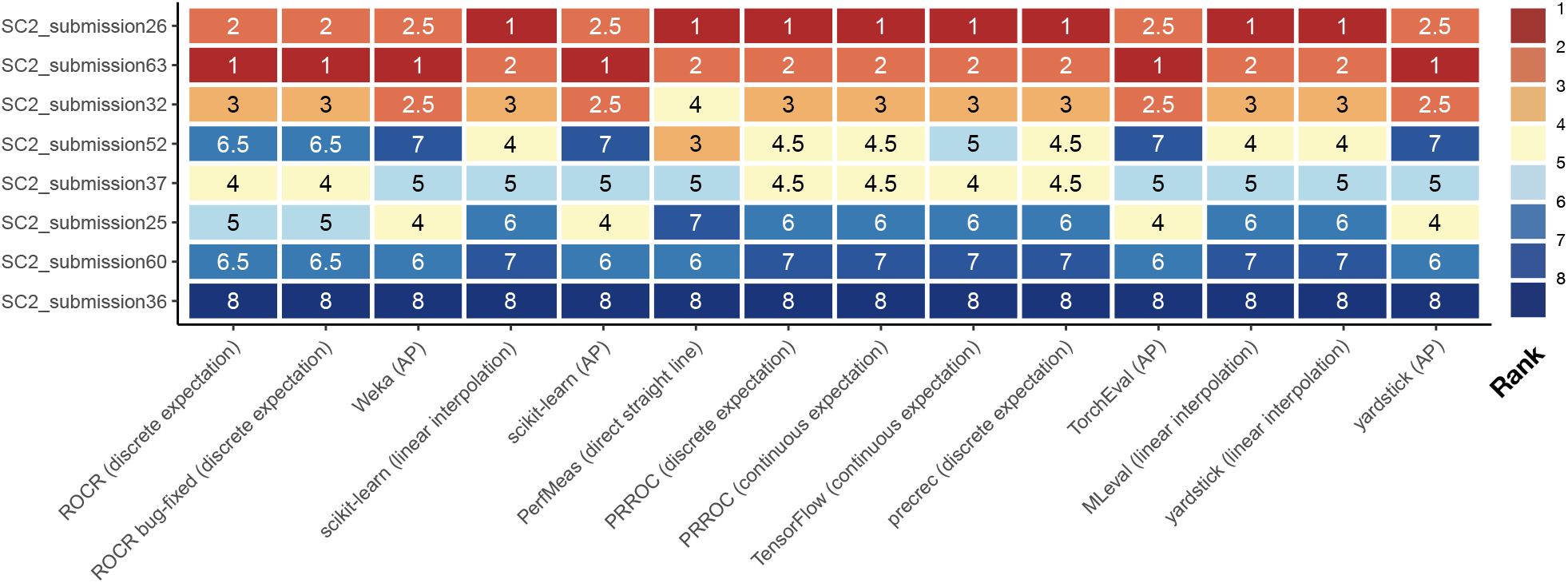
Ranks of the top 8 submissions to the sbv IMPROVER challenge based on the AUPRC values produced by the 10 tools. Each entry shows the rank of a submission based on the corresponding AUPRC value shown in Figure 2b.

**Supplementary Figure 15:**
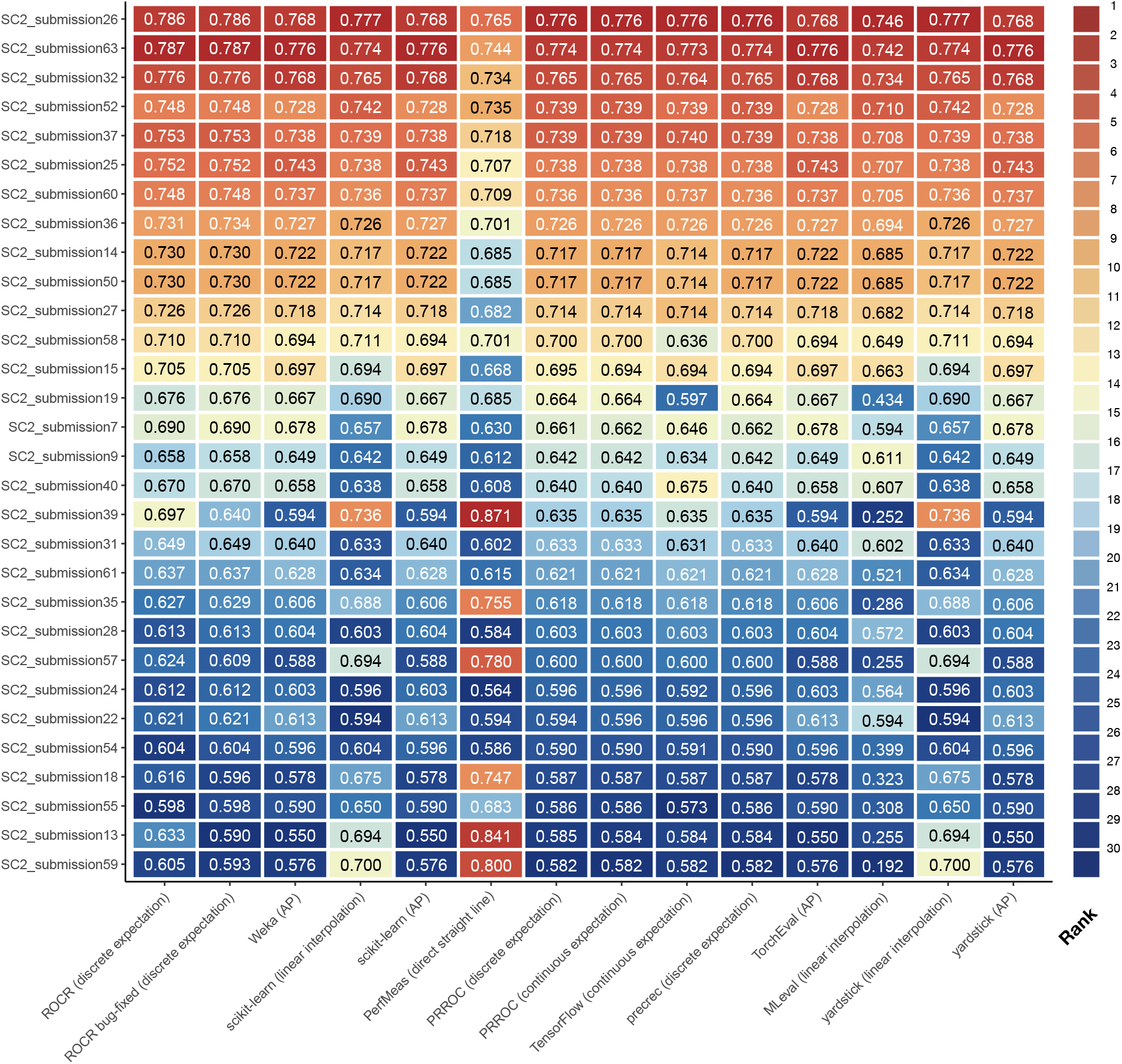
Ranks of the top 30 submissions to the sbv IMPROVER challenge based on the AUPRC values produced by the 10 tools. Each entry shows the AUPRC value and the background color indicates its rank among the submissions.

**Supplementary Figure 16:**
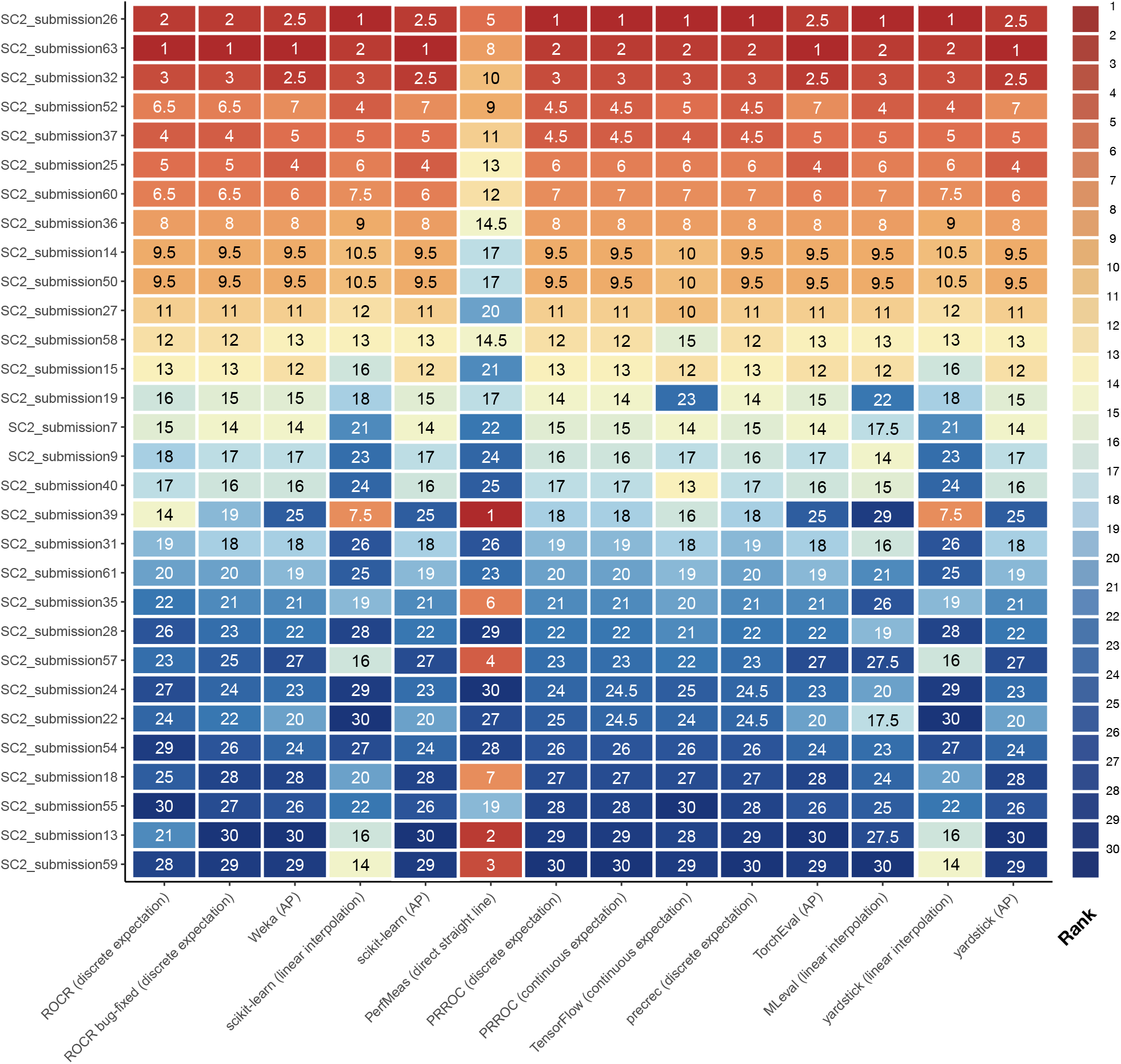
Ranks of the top 30 submissions to the sbv IMPROVER challenge based on the AUPRC values produced by the 10 tools. Each entry shows the rank of a submission based on the corresponding AUPRC value shown in Figure 15.

**Supplementary Figure 17:**
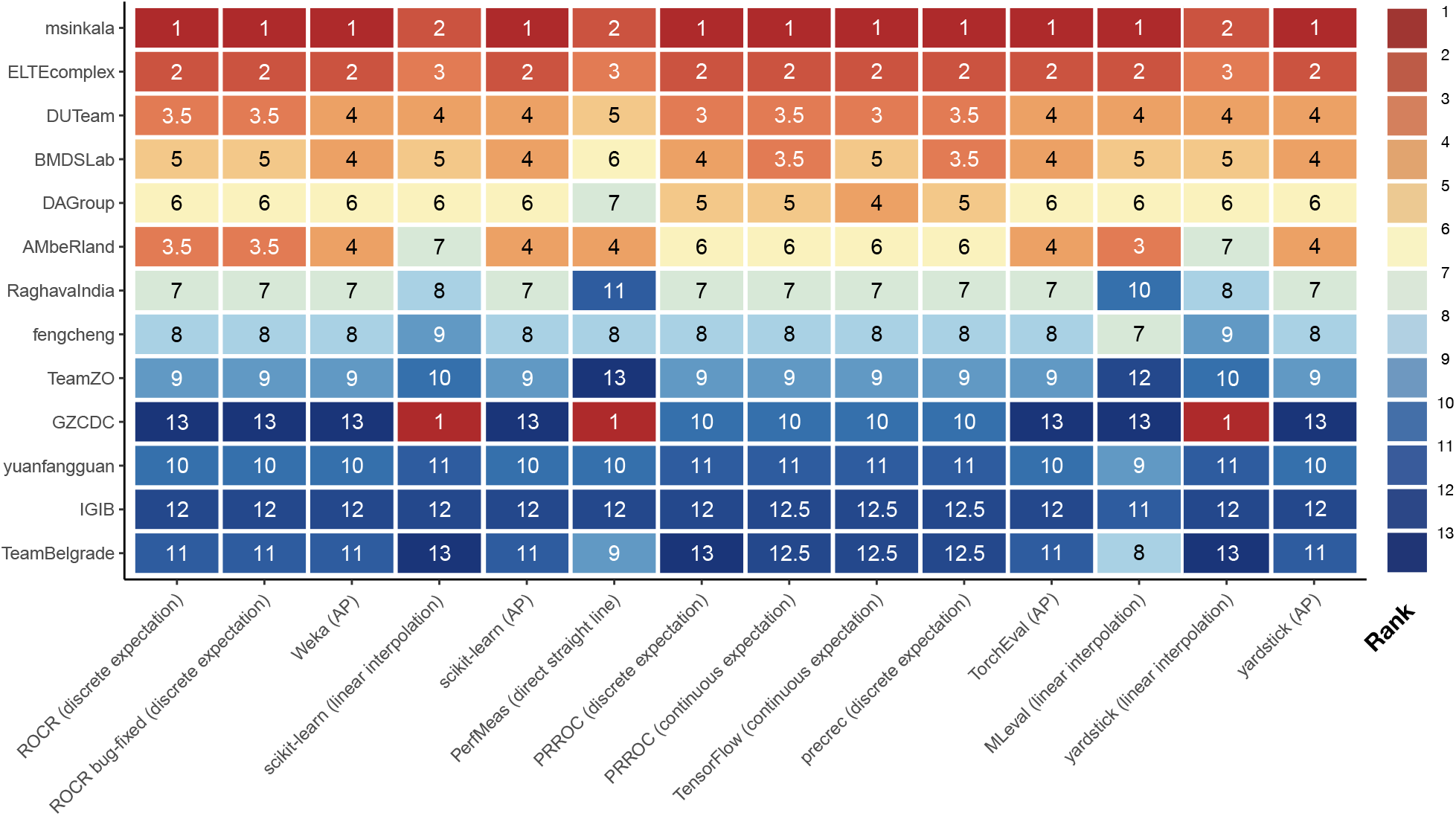
Ranks of the 13 participants of the DREAM Preterm Birth Prediction Challenge based on the AUPRC values produced by the 10 tools. Each entry shows the rank of a submission based on the corresponding AUPRC value shown in Figure 2c.

**Supplementary Figure 18:**
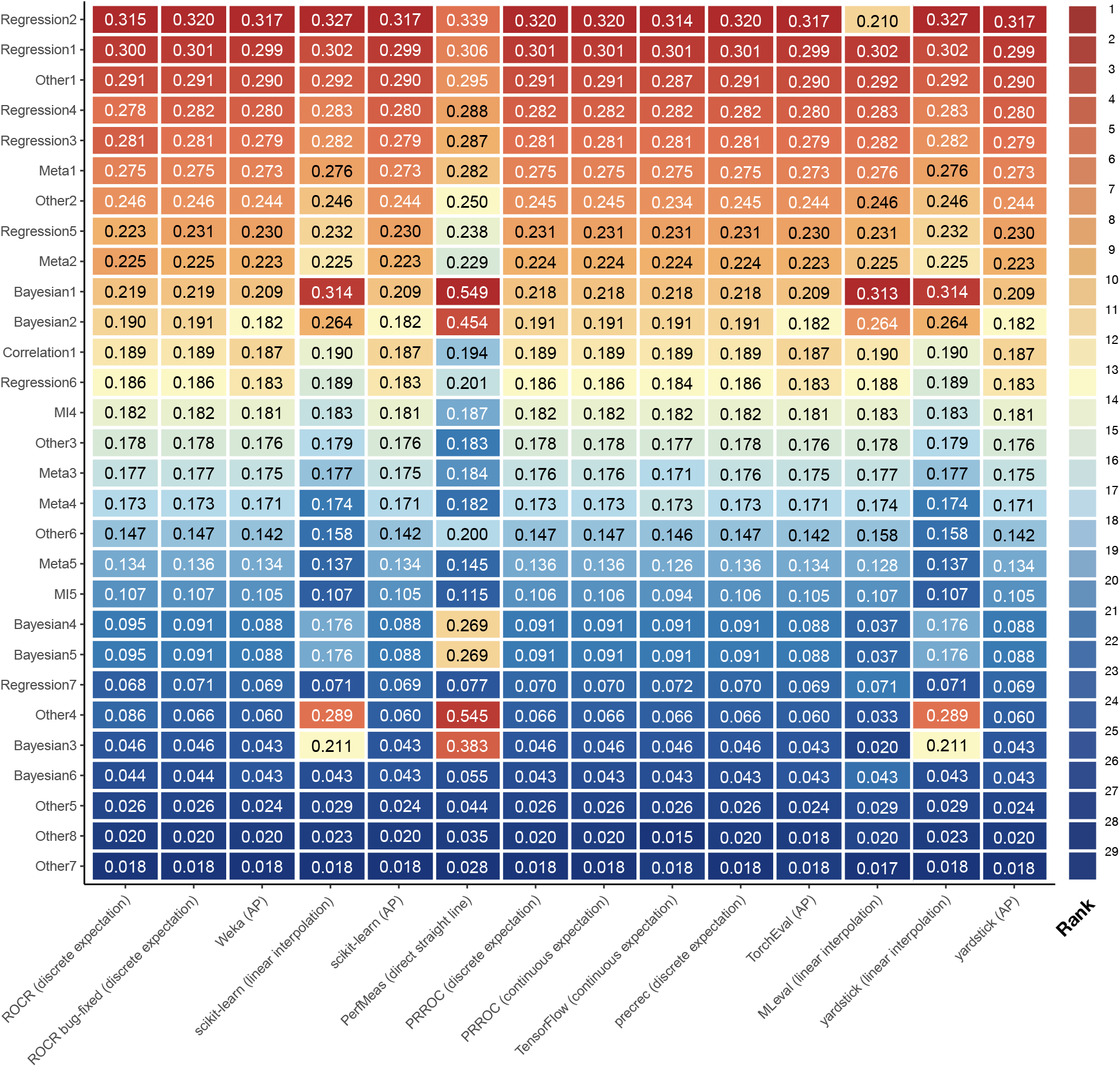
Ranks of the 29 submissions to the DREAM5 transcription factor target challenge based on the AUPRC values produced by the 10 tools. Each entry shows the AUPRC value and the background color indicates its rank among the submissions. Submission “Other8” obtained an AUPRC value of 0.018 from TorchEval (AP), but 0.020 from the other AP method-based tools. This difference was due to TorchEval’s requirement for having the classification scores provided in the form of Tensor objects as input to PyTorch, which led to loss of numeric precision.

**Supplementary Figure 19:**
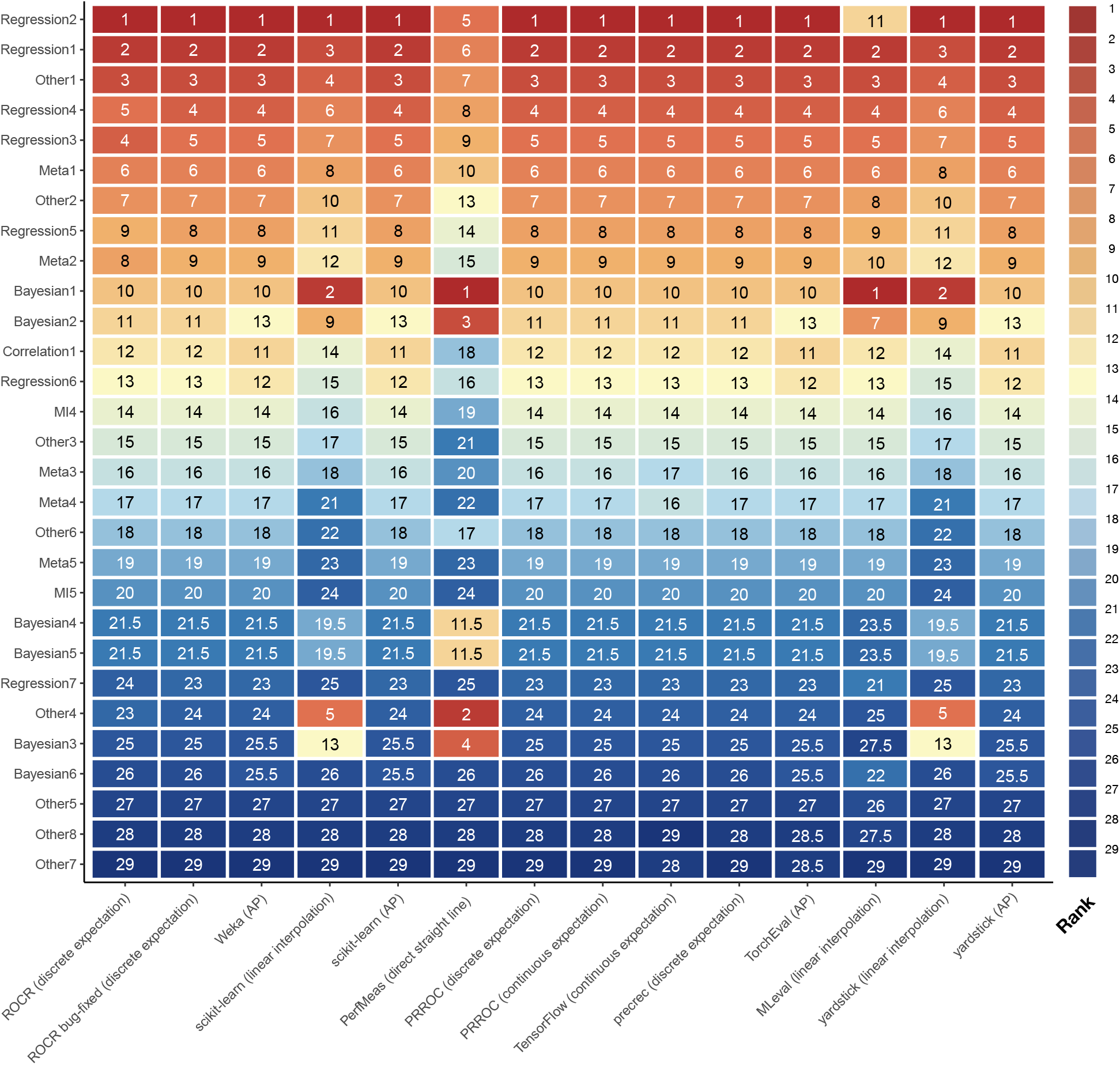
Ranks of the 29 submissions to the DREAM5 transcription factor target challenge based on the AUPRC values produced by the 10 tools. Each entry shows the rank of a submission based on the corresponding AUPRC value shown in Figure 18.

**Supplementary Figure 20:**
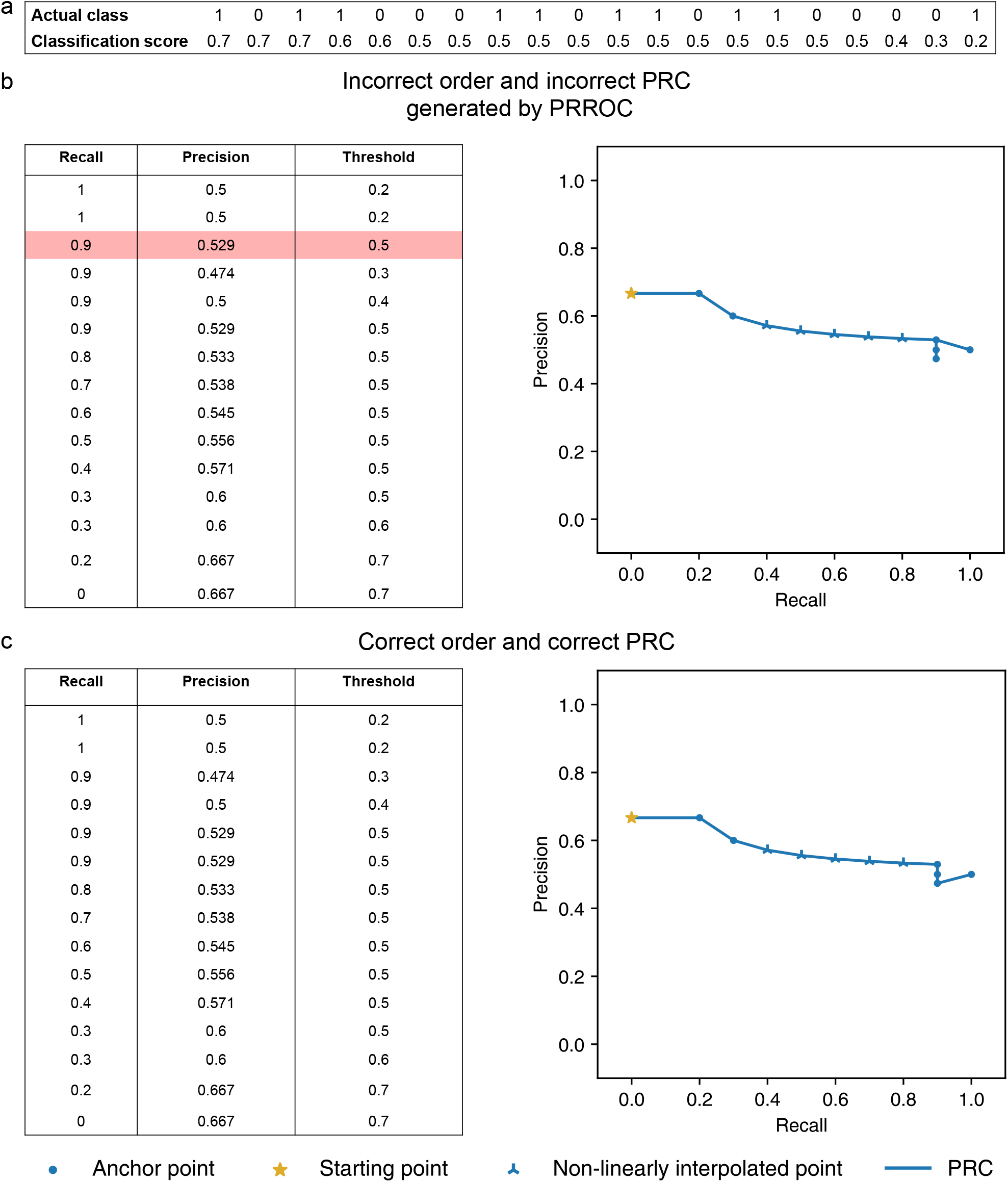
Programming bug of PRROC (option that uses the discrete expectation method to handle ties) when visualizing the PRC. **a** An illustrative data set. **b** Incorrectly ordered thresholds (highlighted in red) produced by the original version of PRROC and the resulting PRC. **c** The correctly ordered threshold and the resulting PRC produced by the bug-fixed version of PRROC.

**Supplementary Figure 21:**
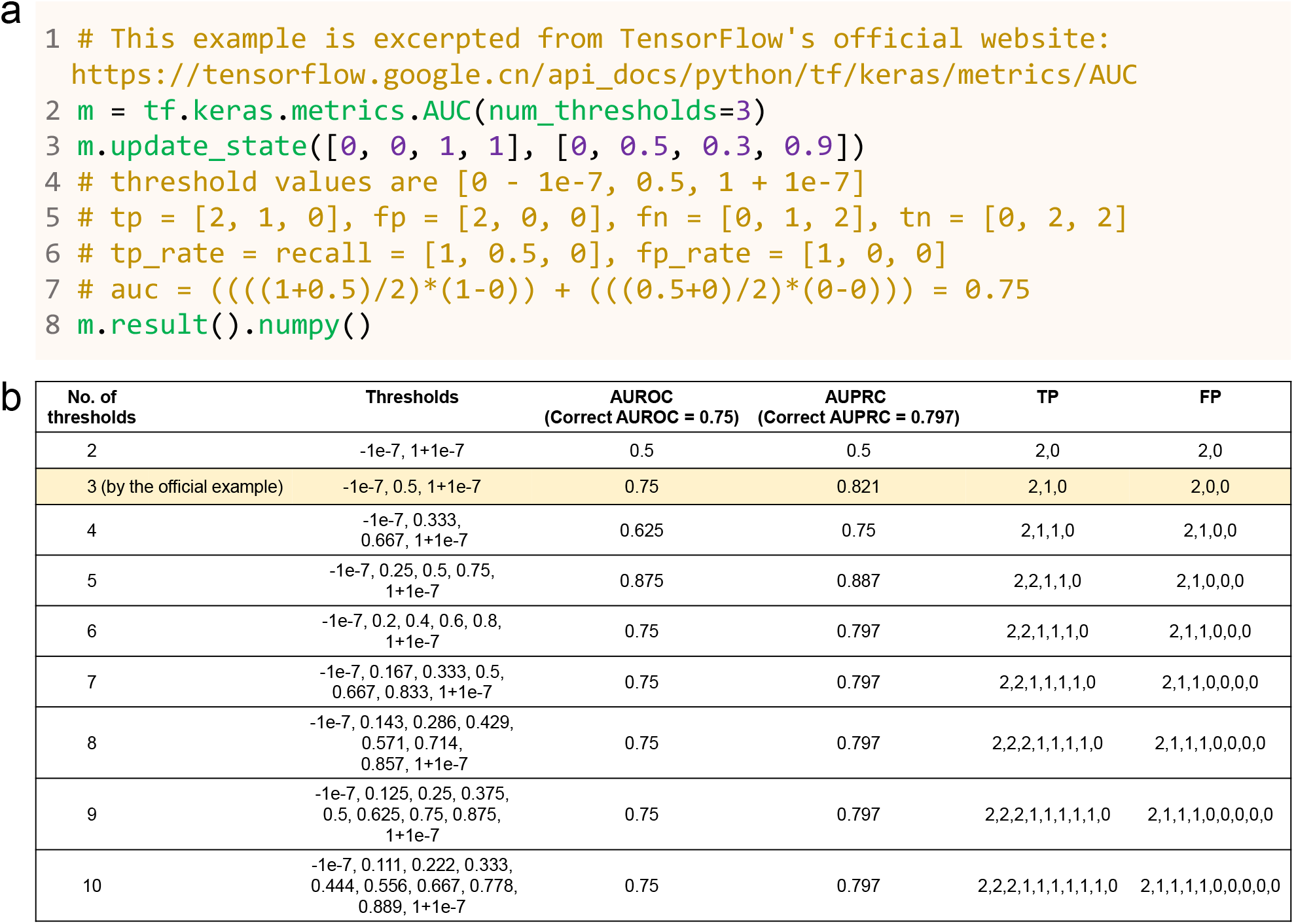
The user-defined number of thresholds can affect TensorFlow’s AUPRC and AUROC calcualtions.**a** An example provided on TensorFlow’s official web site, which uses 3 thresholds. **b** The AUPRC and AUROC values computed by TensorFlow with different numbers of thresholds. The “correct” values, as computed by PRROC using the continuous expectation method to handle ties, are shown in the column headers.

**Supplementary Figure 22:**
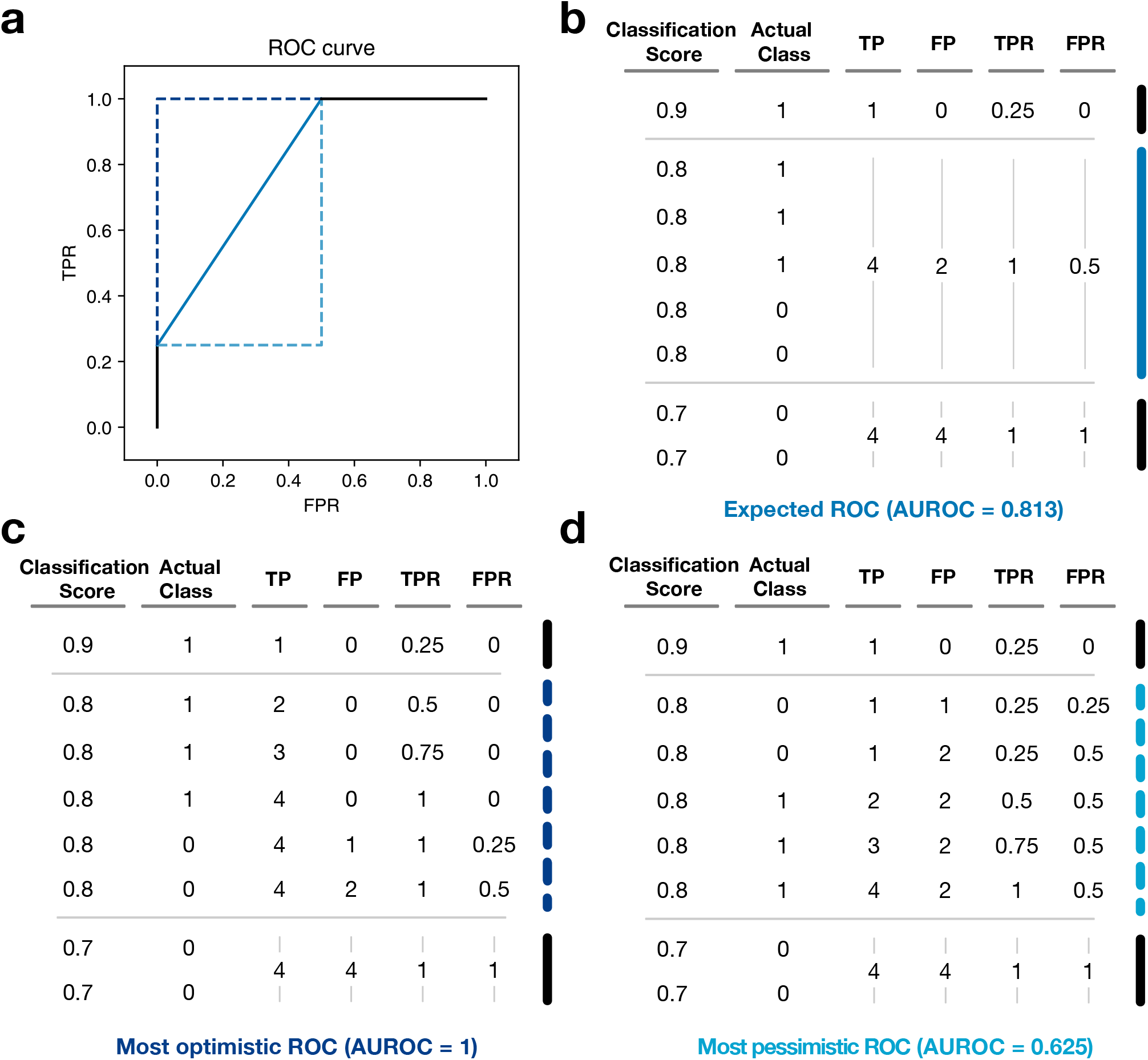
The expected, most optimistic, and most pessimistic ROC curves based on different ways to handle ties in classification scores. **a** The expected ROC curve (the black solid lines and the blue solid line), the most optimistic ROC curve (the black solid lines and the dark blue dashed lines), and most pessimistic ROC curve (the black solid lines and the light blue dashed lines). **b-d** How ties are handled in the three cases, namely taking the average of all possible orders in the expected case (**b**), ordering all actually positive entities first in the most optimistic case (**c**), and ordering all actually negative entities first in the most pessimistic case (**c**).

**Supplementary Figure 23:**
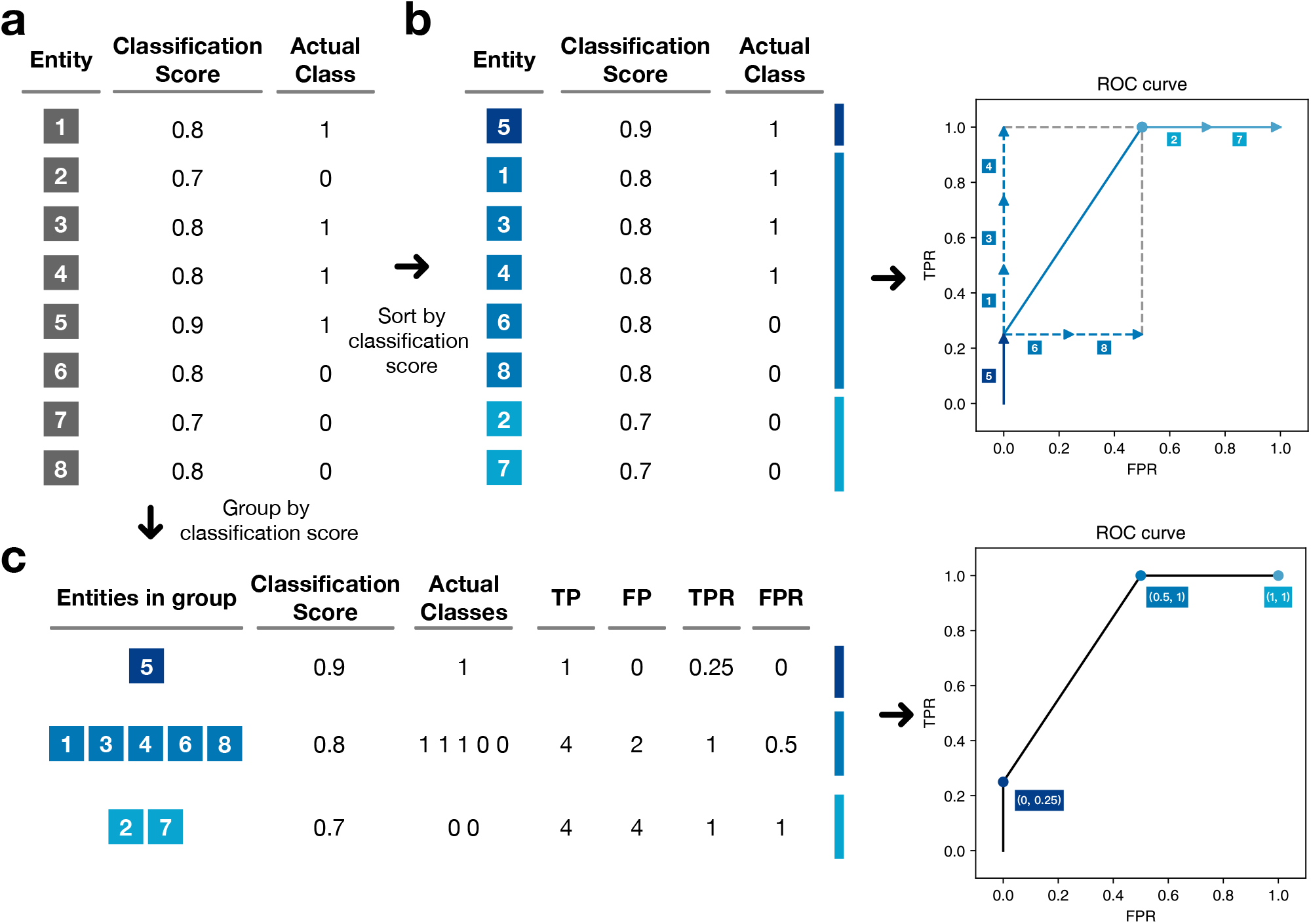
Two algorithms for constructing the ROC curve. **a** An example data set. **b** The workflow of the first algorithm. **c** The workflow of the second algorithm.

**Supplementary Figure 24:**
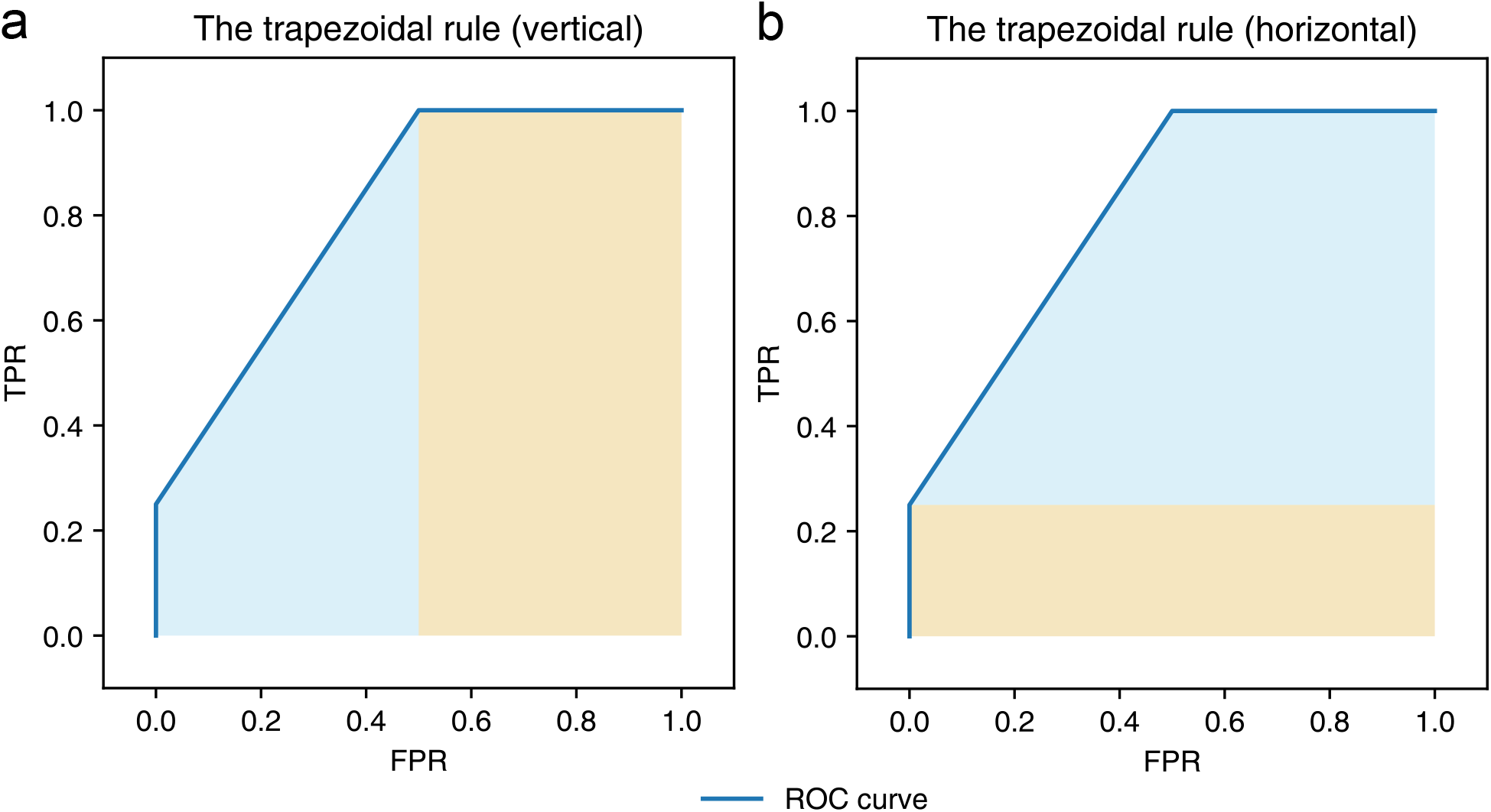
Computing AUROC by the trapezoidal algorithm. The area is divided either vertically **(a)** or horizontally **(b)** before applying the trapezoidal rule. In this example, AUROC is computed by adding up the blue area and the yellow area.

**Supplementary Figure 25:**
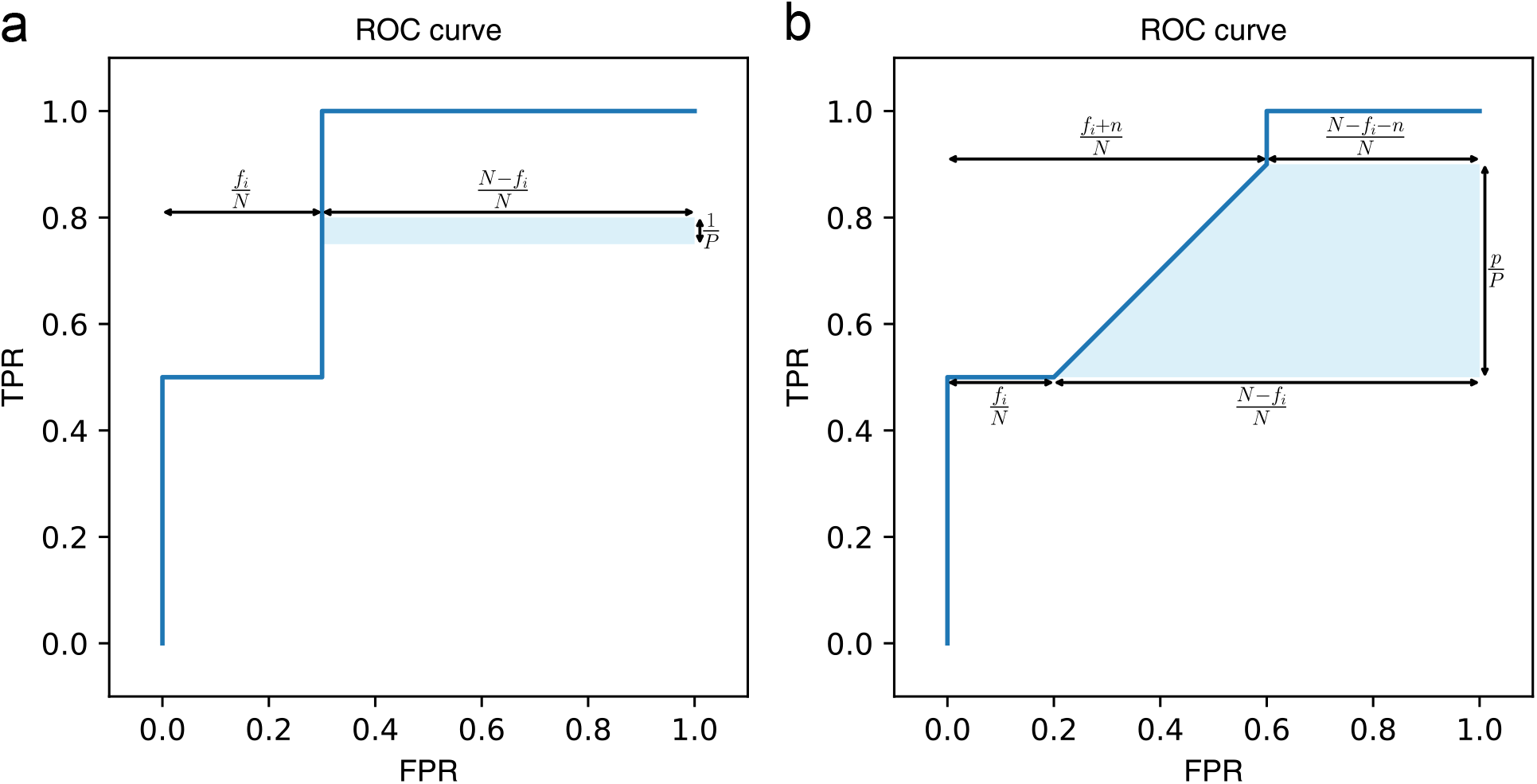
Relationship between the Wilcoxon-Mann-Whitney statistic and AUROC. **a-b** The area gained when there are no ties (**a**) and there are ties (**b**) in the classification scores. All the symbols are defined in the section “Relationship between the Wilcoxon-Mann-Whitney statistic and AUROC” in Supplementary text.

